# SORDINO for Silent, Sensitive, Specific, and Artifact-Resisting fMRI in awake behaving mice

**DOI:** 10.1101/2025.03.10.642406

**Authors:** Martin J. MacKinnon, Sheng Song, Tzu-Hao Harry Chao, Li-Ming Hsu, Scott T. Albert, Yuncong Ma, Tatiana A. Shnitko, Tzu-Wen Winnie Wang, Randy J. Nonneman, Corey D. Freeman, Siddhi S. Ozarkar, Uzay E. Emir, Mark D. Shen, Benjamin D. Philpot, Adam W. Hantman, Sung-Ho Lee, Wei-Tang Chang, Yen-Yu Ian Shih

**Affiliations:** Center for Animal MRI, University of North Carolina at Chapel Hill, Chapel Hill, NC, USA; Biomedical Research Imaging Center, University of North Carolina at Chapel Hill, Chapel Hill, NC, USA; Department of Neurology, University of North Carolina at Chapel Hill, Chapel Hill, NC, USA; Department of Biomedical Engineering, University of North Carolina at Chapel Hill, Chapel Hill, NC, USA; Department of Radiology, University of North Carolina at Chapel Hill, Chapel Hill, NC, USA; Neuroscience Center and Department of Cell Biology & Physiology, University of North Carolina at Chapel Hill, Chapel Hill, NC, USA; Carolina Institute for Developmental Disabilities, University of North Carolina at Chapel Hill, Chapel Hill, NC, USA

**Keywords:** fMRI, tissue oxygen, cerebral blood volume, functional connectivity, brain networks, T1 contrast, rat, mouse, awake, behavior, social, acoustic noise, electromagnetic interference, motion artifacts, susceptibility artifacts, sensitivity, specificity

## Abstract

Blood-oxygenation-level-dependent (BOLD) functional magnetic resonance imaging (fMRI) has revolutionized our understanding of the brain activity landscape, bridging circuit neuroscience in animal models with noninvasive brain mapping in humans. This immensely utilized technique, however, faces challenges such as acoustic noise, electromagnetic interference, motion artifacts, magnetic-field inhomogeneity, and limitations in sensitivity and specificity. Here, we introduce **<u>S</u>**teady-state **<u>O</u>**n-the-**<u>R</u>**amp **<u>D</u>**etection of **<u>IN</u>**duction-decay with **<u>O</u>**versampling (SORDINO), a transformative fMRI technique that addresses these challenges by maintaining a constant total gradient amplitude while acquiring data during continuously changing gradient direction. When benchmarked against conventional fMRI on a 9.4T system, SORDINO is silent, sensitive, specific, and resistant to motion and susceptibility artifacts. SORDINO offers superior compatibility with multimodal experiments and carries novel contrast mechanisms distinct from BOLD. It also enables brain-wide activity and connectivity mapping in awake, behaving mice, overcoming stress- and motion-related confounds that are among the most challenging barriers in current animal fMRI studies.

---

For nearly three decades, gradient-recalled echo (GRE)-based echo-planar imaging (EPI) has been the gold standard in functional magnetic resonance imaging (fMRI), largely due to its ability to rapidly acquire whole-brain volumes with T2* sensitivity to the blood-oxygenation-level-dependent (BOLD) contrast^1–3^ — a broadly utilized marker of brain activity^4–6^. However, despite its widespread use, this technique has several limitations: (1) acoustic noise^7–9^, (2) electromagnetic interference^10–14^; (3) artifacts related to ghosting, motion, and magnetic field inhomogeneity^3,15^; (4) lower sensitivity compared to other neuroimaging modalities^16–18^; and (5) poor spatial specificity due to its bias toward venous vasculature^19,20^. Overcoming these limitations would be groundbreaking for the growing rodent fMRI community^6,21–23^, where key challenges include: (a) acoustic noise from GRE-EPI that forces the use of anesthesia^24,25^, compromising brain function and limiting translational relevance to human fMRI; (b) stress and motion confounds that plague awake rodent fMRI studies unless extensive habituation protocols are implemented^26–33^; and (c) susceptibility artifacts in GRE-EPI that present under the high magnetic field strengths (> 7T) typically used in rodent studies^22,34,35^.

“Zero” acquisition delay imaging sequences, such as zero echo time (ZTE), RUFIS, PETRA, and MB-SWIFT, primarily known for structural MRI applications^36–38^, acquire data just a few microseconds after RF pulsing and offer potential solutions to some of the challenges faced in fMRI. In these sequences, imaging gradients ramp to a plateau, and center-out frequency-encoding spokes capture the free-induction-decay (FID) induced by RF pulses under steady gradients. The data from multiple radially oriented spokes are then reconstructed to form images. With proper encoding trajectories and the ZTE feature, these sequences are inherently less susceptible to acquisition-related acoustic noise, electromagnetic interference, and artifacts^36,37^. Although some of these zero acquisition delay techniques have been adapted for fMRI^39–43^, they still face a critical limitation: none have demonstrated robust sensitivity compared to the gold standard GRE-EPI. This lack of sensitivity hinders their broader adoption, as the compromised sensitivity requires compensation with more experimental repetitions or larger sample sizes. Additionally, due to their extremely short acquisition delays, these sequences are inherently insensitive to T2- or T2*-weighted BOLD contrast, highlighting the need to better understand alternative fMRI contrast mechanisms for functional neuroimaging.

In this study, we address several major barriers in fMRI, including acoustic noise, electromagnetic interference, ghosting, motion, susceptibility artifacts, sensitivity, and specificity by introducing a novel fMRI technique: **<u>S</u>**teady-state **<u>O</u>**n-the-**<u>R</u>**amp **<u>D</u>**etection of **<u>IN</u>**duction-decay with **<u>O</u>**versampling (SORDINO), named for its analogy to the muting of a musical instrument. Unlike existing ZTE techniques acquire data only on steady gradients, SORDINO takes a fundamentally different approach and further improves the benefits of zero-acquisition-delay sequences: it maintains constant gradient amplitudes throughout the entire sequence while continuously changing the gradient direction, providing minimal acoustic noise and gradient-related artifacts and maximal sensitivity by acquiring data exclusively during the gradient ramps. We benchmarked SORDINO against GRE-EPI and ZTE on a high-field preclinical MRI system (9.4T), demonstrated SORDINO’s exceptional compatibility with simultaneous electrophysiology, electrochemistry or calcium imaging at cellular resolution, modeled and revealed its contrast mechanisms for functional neuroimaging, and showcased its ability to measure brain-wide resting-state connectivity in awake mice. Furthermore, we demonstrated that SORDINO facilitates experiments that are otherwise highly challenging, if not impossible, including mapping naturalistic circuit activity in behaving, head-fixed mice and simultaneously imaging two mouse brains during social interactions. Our findings suggest that SORDINO offers robust sensitivity and is a transformative technique for functional brain mapping, particularly suited for subjects that cannot tolerate acoustic noise or are susceptible to magnetic field inhomogeneity and motion artifacts. Through this work, we release the SORDINO sequence and its corresponding database, providing the field with an accessible resource to enable new research opportunities and further technical advances.

## RESULTS

### SORDINO sequence

To date, no MRI technology has achieved imaging by clamping both total gradient amplitude (G_TotalAmp_) and gradient angular change (α) throughout the entire duration of a scan. Our creation of SORDINO started with one very simple thought: MRI scans produce loud acoustic noise due to vibrations in the gradient coils driven by rapidly switching electrical currents, and this is also the source of many Eddy-current-related artifacts, including ghosting and distortion in EPI. Thus, the question arises: How can we efficiently distribute the gradient changes needed for a 3D encoding without ever resetting the gradient? Following this thought, we identified the simplest strategy to achieve 3D encoding with the lowest possible α. The sequence requires a one-time initialization at the start of an fMRI experiment, as depicted in **Figure 1a & Figure S1a**. Following this initialization, both G_TotalAmp_ and α remain constant throughout the entire acquisition (i.e., dG_TotalAmp_/dt=0 and dα/dt=0). This condition is held not only across all encoding datapoints within an imaging volume, but also across all volumes (i.e., all time points) during an fMRI experiment. An analogy for SORDINO’s gradient control to existing radial sampling methods is akin to the smooth, continuous motion of a modern clock’s second hand versus the discrete, ticking motion of a traditional clock. While the gradient direction constantly changes, we apply a phase-incrementing, spatially non-selective RF pulse to induce the FID signal while reducing spurious echoes (**Figure S1b**). Given the presence of the gradient field, the FID naturally realizes “frequency encoding” in the form of a curved, center-out k-space line (i.e., a spoke). Acquiring data during the ramping gradient incurs no noticeable penalty (**Figure S1c&d**). **Figure 1b** provides a conceptual representation of the spoke evolution in the 2D plane, illustrating that the end of each spoke points towards the beginning of the next spoke. **Figure 1c** & **Video 1** visually demonstrate the precise gradient trajectory employed in SORDINO, where the last spoke of one volume aligns seamlessly with the first spoke of the next volume. This strategy ensures a constant, minimal change in gradient direction, eliminating the need for resets. This straightforward approach results in an ultra-low slew rate for fMRI (0.21 T/m/s), which is four orders of magnitude lower than EPI (1263.62 T/m/s) and nearly one hundred times lower than other 3D zero-acquisition-delay sequences such as ZTE or MB-SWIFT (15.66 T/m/s) at identical spatiotemporal resolution (see Methods for parameter details). Furthermore, by eliminating traditional gradient ramping and settling times (**Figure S1a**) while incorporating an oversampling anti-aliasing strategy (**Figure S1e&f**), SORDINO can achieve a lower bandwidth and maximize acquisition time, which translates directly to improved signal-to-noise ratio (SNR) and having fewer datapoints that fall within the T-R switch dead time (**Figure S1g-i**) compared to other short acquisition delay fMRI methods. Further details for the SORDINO sequence are presented in Methods.

**Figure 1.**
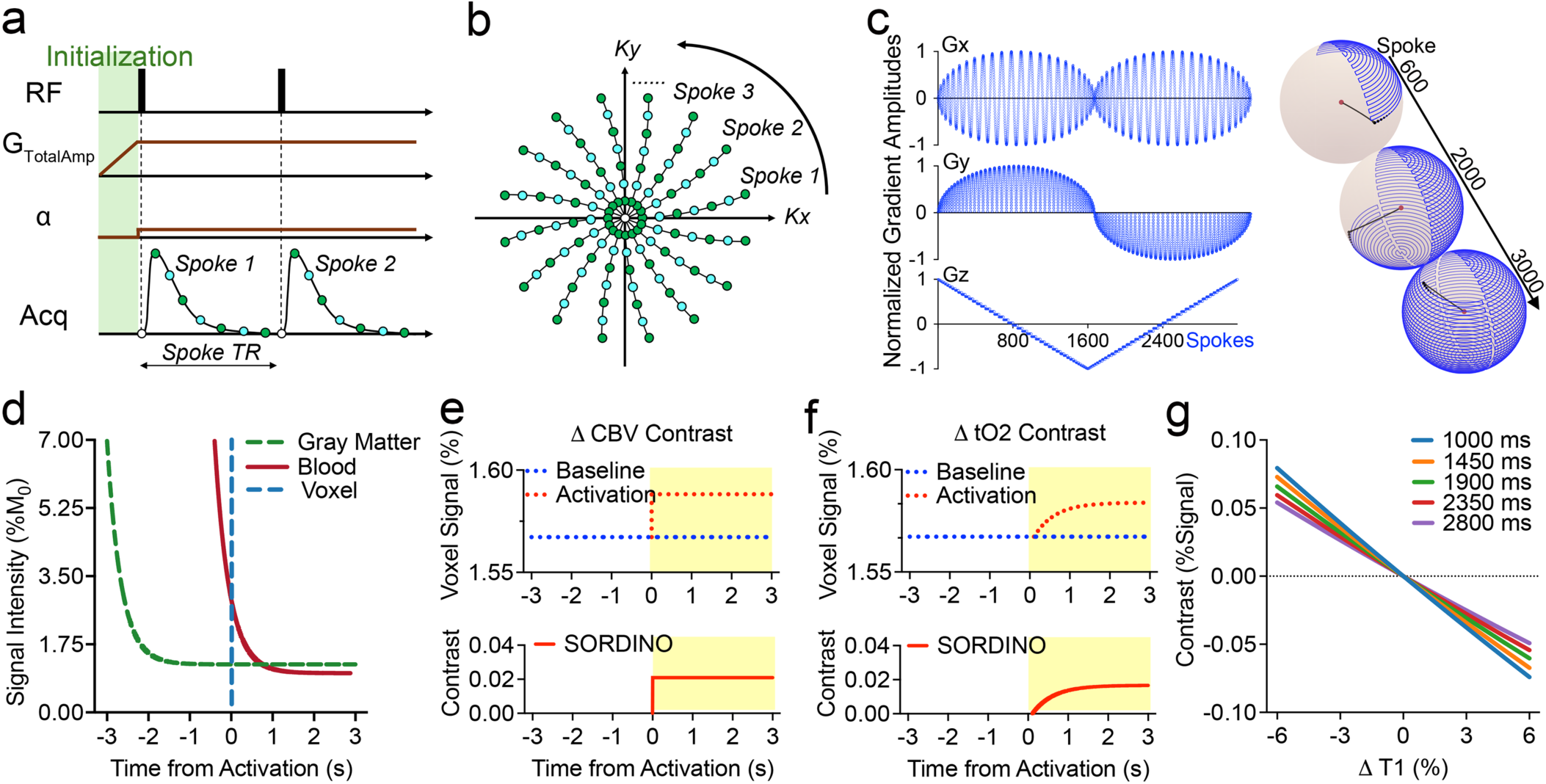
SORDINO sequence and modeling of its functional contrast. **(a)** In SORDINO, the gradient changes constantly at an angular change rate α and has an extremely low slew rate, thus eliminating acoustic noise and Eddy current artifacts. Data acquisition commences exclusively and continuously throughout the non-stop gradient ramping, thus enhancing SNR. In this conceptual plot, dark green datapoints represent acquisition at nominal bandwidth and light blue datapoints represent the oversampled digitized datapoints. **(b)** Conceptual spoke evolution in a 2D plane, showing multiple curved spokes in k-space. **(c)** SORDINO gradient trajectory in 3D. **(d)** In the case of using a head-only excitation coil, the measured voxel signal is comprised of a mixture of stationary tissue and transient blood signal. Bloch equation simulations show that magnetization of stationary tissue reaches a steady-state within a few seconds (green dashed line), whereas the blood signal (red line) does not reach a steady-state at the time of measurement due to the short blood transit time, as fresh arterial blood would experience fewer RF excitations. Specific parameters for these simulations are detailed in Methods. **(e)** CBV contributions at the voxel level, considering: arterial transit time = 280 ms, baseline blood flow = 10 mm/s, local flow acceleration = 1.5 mm/s, activation radius = 1 mm, blood/tissue volume fraction = 5%, and CBV increase = 20%. Regionally accelerated blood would be subjected to 13 fewer RF-pulses and heightened the blood signal, resulting in a robust increase in SORDINO signal. **(f)** tO_2_ contribution at the voxel level: a physiological 30 µM transient increase in tissue oxygen corresponds to a 32 ms decrease in T1 from a baseline of 1900 ms (oxygen T1 relaxivity is 0.3 mM^-1^s^-1^.) The tO_2_-induced T1-shortening results in a measurable increase in the SORDINO-fMRI signal, taking approximately 1 s to establish a new steady-state. **(g)** The relationship between T1 changes and SORDINO-fMRI contrast is nearly linear across a range of gray matter T1 values (see Methods for detailed calculations).

### SORDINO Contrast Modeling

First, we conducted a series of modeling experiments to explore a range of possible parameters capable of producing contrasts in response to T1 changes. These models served as a foundation as we examined the impact of specific signal modulators within the rodent brain at 9.4T: inflow-enhanced cerebral blood volume (CBV) and alterations in tissue oxygen levels. Conventionally, the inflow effect observed in fMRI is primarily attributed to the apparent T1 shortening resulting from incoming spins that experience few or no RF pulses^44,45^, which generates a stronger signal compared to that from stationary tissue^44^. When using SORDINO with head-only RF excitation, frequent RF pulsing rapidly leads the magnetization of stationary tissue to a steady-state (**Figure 1d**, green dashed line); however, inflowing arterial blood experiences fewer RF pulses, resulting in a higher signal compared to static tissue (**Figure 1d**, red line and blue vertical dash line). Collectively, the combined voxel signal stabilizes at a steady-state when the blood transit time and CBV remain stable (**Figure 1d-f**, blue dashed lines). We subsequently modeled the effect of activation-induced increases in CBV. Given that the blood signal is higher than the tissue (see **Figure 1d**, blue vertical dash line), an increase in vascular-space-occupancy^46,47^ translates into positive changes in the SORDINO signal (**Figure 1e**). **Figure S2a & b** display a series of raw SORDINO images of a rat brain in live and postmortem conditions, respectively. These images also feature the robust CBV contrast in the live condition as predicted when using a head-only coil for RF excitation. Raw SORDINO images across distinct spatial resolutions are shown in **Figure S2c**.

Next, we evaluated the influence of alterations in tissue oxygen (tO_2_) levels. Guided by invasive tO_2_ ground-truth measurements^48^ and prior studies revealing the relationship between molecular oxygen concentration and T1 relaxation rate (R1)^49^. We estimated that changes in tO_2_ contribute to SORDINO contrast to a similar order of magnitude as CBV (**Figure 1f**). Given the well-documented spatial specificity of tO_2_ and CBV metrics^50–52^, SORDINO emerges as a compelling alternative to GRE-EPI BOLD-fMRI (*vide infra*). Furthermore, it’s worth noting that SORDINO contrast exhibits linearity across a spectrum of T1 values in both the rodent brain at 9.4T and the human brain at 3T, and this characteristic positions SORDINO to accurately map brain activity across the physiological range of T1 changes (**Figure 1g**). In addition to using a head-only coil for RF excitation, we explored SORDINO contrast when employing a whole-body coil for RF excitation. Our findings indicate that, in this scenario, tO_2_ becomes the dominant contributor to SORDINO contrast, while CBV increases result in slight negative changes due to the longer T1 of blood compared to brain tissue (**Figure S3**).

### SORDINO Acquisition Parameters and Sampling Efficiency

To examine our operational hypothesis regarding the use of SORDINO for fMRI, it is imperative to model the efficacy of SORDINO parameter selection in delineating T1-related functional activation and to validate these parameters *in vivo* (**Figure S4**). In **Figure S4a-c**, we display various combinations of repetition time (TR) and flip angle (FA). While a shorter spoke-TR constrains T1 recovery, it allows more spokes to be acquired within a fixed volume-TR (**Figure S4d**). We thus considered an efficiency factor, calculated as the square root of spoke-TR^53^, to account for the influence of the number of spokes acquired within a fixed volume-TR. In **Figure S4e-f**, we present the outcomes of Bloch equation modeling, which estimates the efficiency of SORDINO acquisition. Our modeling reveals that different TR-FA combinations yield varying levels of sensitivity to T1 changes, and that FA ranging from 2° to 5° offer optimal efficiency when maintaining a spoke-TR between 0.6 and 2.4 ms (**Figure S4g**). To validate these modeling results, we conducted experiments using a forepaw stimulation task in rats, assessing five different TR-FA combinations. The detected functional activations *in vivo* (**Figure S4h**), as determined through contrast-to-noise ratio (CNR) analysis, qualitatively aligns with the modeled SORDINO imaging efficiency (**Figure S4g**).

### SORDINO versus GRE-EPI

Next, we benchmarked SORDINO against the gold-standard GRE-EPI across various metrics, including acoustic noise, electromagnetic interference, ghosting, distortion, susceptibility artifacts, motion artifacts, sensitivity, and specificity. To ensure a fair comparison, we configured the SORDINO and EPI parameters to achieve identical spatial and temporal resolution (see Methods for detailed parameter settings).

#### Negligible Acoustic Noise

SORDINO demonstrated exceptional proficiency in mitigating acoustic noise, as illustrated in **Figure 2a**. Across the recorded frequency range (0-15 kHz), the sound pressure level (SPL) profile for SORDINO closely follows the scanner’s idle state. In comparison to SORDINO, the SPL is up to 20 dB higher in EPI and 5 dB higher in ZTE. This acoustic noise reduction in SORDINO holds the potential for significant benefits in awake animal fMRI, where it can alleviate subject stress and reduce habituation time (**Figure 2b&c**). Specifically, in our experimental conditions, we found that head-fixed awake mice subjected to longitudinal SORDINO experiments exhibited elevated cortisol levels only on the first day of imaging, with cortisol level decreasing on Day 3 and showing no significant deviation from the baseline on Day 5. This stands in stark contrast to awake mice undergoing GRE-EPI scans, which consistently displayed heightened cortisol levels throughout the measured time points (**Figure 2b**). This reduction in stress was also evident in the subjects’ body weight changes over time (**Figure 2c**).

**Figure 2.**
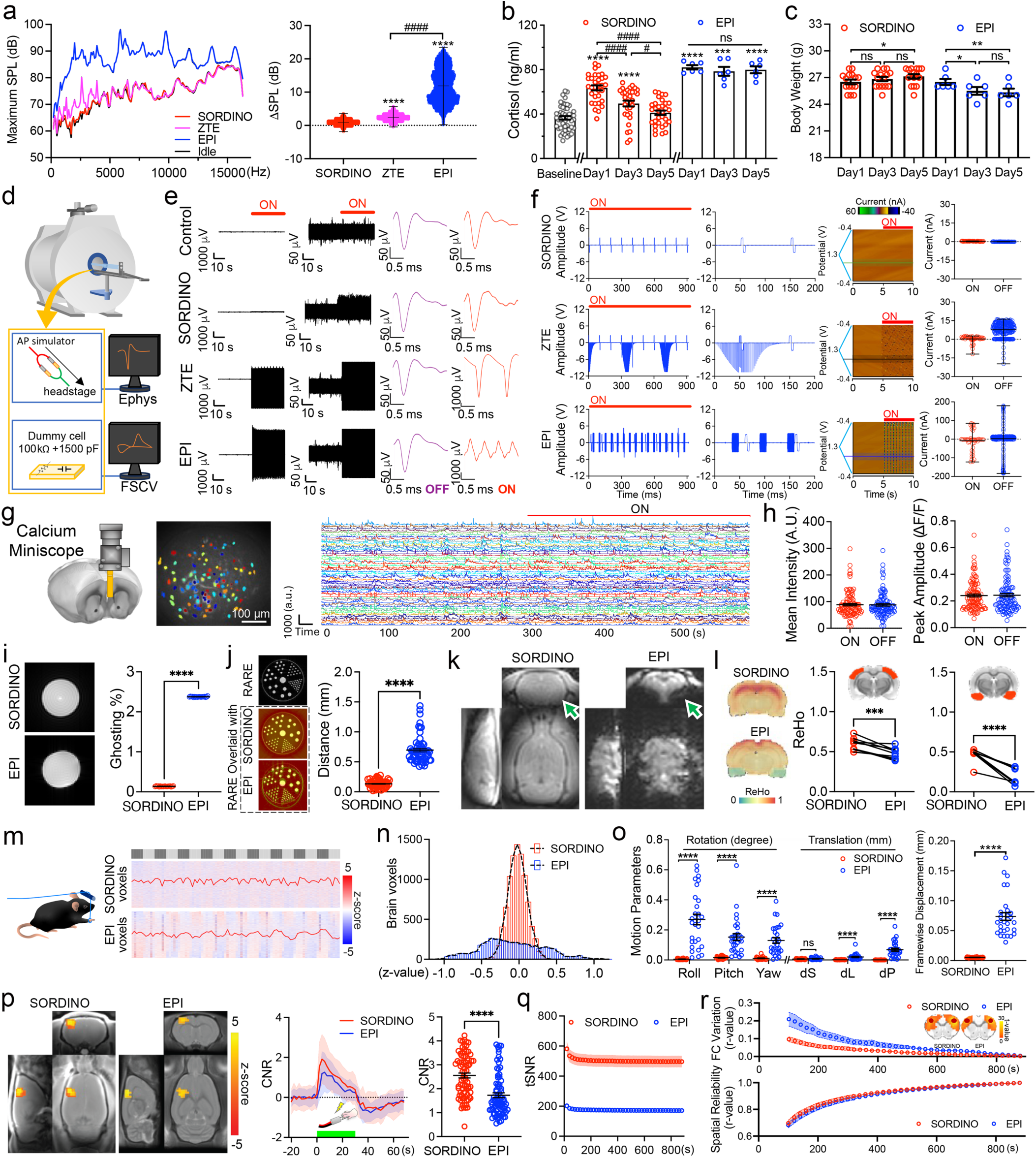
SORDINO versus EPI. **(a)** Comparison of acoustic noise induced by active SORDINO, ZTE, and EPI scanning against the scanner idle state. SORDINO effectively eliminates gradient-related acoustic noise, enabling silent imaging (n = 10 trials; one-way ANOVA, F(2,1499) = 20694, *p*-values are from Tukey’s HSD). **(b, c)** Mice undergoing a five-day head-fixation habituation process exhibit reduced stress hormone levels (Serum Cortisol: n = 49 baseline, n =32 SORDINO and n = 6 EPI; One-Way ANOVA;, F(6,156) = 38.79, *p* < 0.0001; *p*-values are from Tukey’s HSD) and maintain body weight in the SORDINO group but not in the EPI group (n = 15 SORDINO; Repeated Measures one-way ANOVA followed by Bonferroni multiple comparison test, effect of MRI sequence, F(1.87,26.10) = 6.76, *p* = 0.005; n = 6 EPI; Repeated Measures one-way ANOVA followed by Bonferroni multiple comparison test, effect of MRI sequence, F(1.45,7.26) = 12.65, *p* = 0.006. **(d)** Spike simulator and dummy cell setup for quantifying electromagnetic interference. **(e)** Electrophysiological recordings in MRI with and without active SORDINO, ZTE, and EPI scanning show that SORDINO significantly reduces electromagnetic interference, enabling online sorting of simulated neuronal spiking signals without additional pre-processing. **(f)** Electrochemical recording using fast-scan cyclic voltammetry in MRI demonstrates that SORDINO allows real-time recording without requiring the interleaved recording approach previously reported for EPI. Results are presented as the mean oxidation current recorded at 0.65V vs. Ag/AgCl reference electrode with the corresponding range. **(g)** SORDINO’s versatility supports the placement of a calcium miniscope above the mouse head, enabling unprecedented real-time calcium imaging at cellular resolution during brain-wide fMRI. Calcium activities from 103 neurons in the prelimbic cortex (PrL) are displayed during SORDINO off and on periods. **(h)** Active SORDINO scanning does not compromise calcium imaging quality, as indicated by consistent mean intensity and peak amplitude measurements. **(i)** Ghosting artifacts are absent in SORDINO compared to EPI (n = 19 independent trials; two-tailed paired t-test, t(18) = 1006.18, *p* < 0.0001). **(j)** Using Cartesian-sampled spin-echo data as the ground truth, SORDINO shows no structural distortion compared to EPI (n = 60 measurements; two-tailed paired t-test, t(59) = 21.26, *p* < 0.0001). **(k)** In vivo comparison of SORDINO and EPI images from the same subject under identical shimming reveals reduced susceptibility artifacts in several regions, including the amygdala (arrowheads). **(l)** Regional homogeneity (ReHo) analysis on resting-state data indicates preserved local correlation in susceptibility artifact-prone regions with SORDINO, while EPI shows nearly zero ReHo in these regions (n = 9; two-tailed paired t-test, t(8) = 7.49, *p* = 0.0003). **(m)** Passive forelimb movement created by using a box-design in a deeply anesthetized mouse demonstrates that SORDINO drastically reduces motion-correlation artifacts. **(n)** Correlation histograms depict brain voxels with strong positive and negative correlations to the motion paradigm in EPI, which are absent in SORDINO. **(o)** Six motion parameters and FD extracted from resting-state SORDINO and EPI acquisitions show that SORDINO is immune to motion confounds (n = 30 subjects; two-tailed paired t-test between all parameters: t(29) = 1.36 – 7.60, *p* < = 0.0000–0.19; Two-tailed paired t-test between groups in FD: t(29) = 10.53, *p* < 0.0001). **(p)** SORDINO demonstrates robust sensitivity compared to GRE-EPI-BOLD using a within-subject, within-session design at identical spatiotemporal resolution (n = 16 subjects, 80 unique trials; linear mixed effects model, β = 0.82, *p* < 0.0001). **(q)** tSNR in the somatosensory cortex comparison from resting-state SORDINO data (n = 9). The higher tSNR in SORDINO allows the rather small T1-related changes to be detected. **(r)** SORDINO demonstrates comparable resting-state FC measurement to GRE-EPI-BOLD. FC variation and spatial reliability are measured using seed-based analysis, leveraging the intrinsic homotopic FC feature in the somatosensory cortex across hemispheres. Randomly sampled time-series data lengths (100 random selections per length) highlight the comparable time needed to depict reliable FC compared to the full data length. Results are expressed as mean ± SEM. \**p*<0.05, \*\**p*<0.01, \*\*\**p*<0.001 and \*\*\*\**p*<0.0001 compare with SORDINO, baseline, or indicated groups; #*p*<0.05 and ####*p*<0.0001 compare with indicated groups. ns, not significant.

#### Ultra-low Electromagnetic Interference

We performed electrophysiological and electrochemical recordings (fast-scan cyclic voltammetry, FSCV)^48^ during active MRI scanning. These experiments were performed *in vitro*, allowing the measurement of repeatable ground-truth signals generated by a spike signal simulator without confounders such as physiological noise (**Figure 2d-f**). Remarkably, due to the absence of transient gradient switching, SORDINO exhibited minimal gradient-induced artifacts in spike recording, which did not hinder the ability to identify spikes using a conventional online sorting approach (**Figure 2e**). Additionally, there was no noticeable difference in FSCV readout between SORDINO off and on states (**Figure 2f**). These are in sharp contrast to GRE-EPI and ZTE, where the induced artifacts have long been a barrier to continuous and real-time measurement of these electrical based physiological signals during fMRI. This drastically reduced electromagnetic interference also enables the new capability of acquiring calcium imaging data at cellular resolution using a head-mounted miniscope during simultaneous fMRI (**Figure 2g** & **Video 2**). Analysis of calcium signal dynamics during SORDINO off and on periods showed no significant differences in calcium imaging quality (**Figure 2h**).

#### Minimized Ghosting, Distortion, Susceptibility, and Motion Artifacts

Using an established analysis approach^54^, we showed the lack of ghosting in SORDINO (**Figure 2i**). We also assessed spatial distortion by comparing data against Cartesian-sampled turbo spin echo (TurboRARE) images, showing the lack of distortion in SORDINO compared to EPI (**Figure 2j**). To compare susceptibility artifacts in SORDINO and EPI, we presented raw images in **Figure 2k**, highlighting areas that are typically most prone to air-tissue interface dephasing and signal drop out in EPI, which are preserved in SORDINO. To further evaluate the impact of susceptibility artifacts, we analyzed resting-state data to generate regional homogeneity (ReHo) images^55^, demonstrating measurable ReHo in regions that are challenging to assess by EPI (**Figure 2l**). In addition, we remotely induced passive limb movements in deeply anesthetized mice by pulling a nylon monofilament string (**Figure 2m** & **Video 3**) and demonstrated a marked reduction in unwanted motion-induced correlation in SORDINO compared to EPI (**Figure 2n**). Further, we performed SORDINO and EPI scans in two groups of awake mice and extracted motion parameters from the time-series data. Our results demonstrated a significant decrease in motion parameters and framewise displacement (FD) using SORDINO (**Figure 2o**).

#### Robust Sensitivity

Using an established forepaw electrical stimulation task in rats, we assessed the contrast-to-noise-ratio (CNR) against baseline standard deviation and presented data in z-scores. SORDINO consistently demonstrated a robust sensitivity to functional activation within the primary somatosensory cortex (S1) of the forepaw region. The CNR was comparable, if not superior, to that of EPI (**Figure 2p**). In another cohort of subjects, we also compared forepaw-evoked responses obtained from SORDINO and GRE-EPI under different conditions and acquisition parameters, demonstrating comparable CNR (**Figure S5**). Further, our analysis of time series data from the rat brain revealed a higher tSNR with SORDINO (**Figure 2q**). Moreover, we benchmarked the efficiency of SORDINO for resting-state FC measurement by performing seed-based analysis to calculate FC variation and spatial reliability of the somatosensory homotopic FC pattern. These results, presented as a function of randomly sampled time-series data lengths (100 random selections for each length to compute the mean and standard deviation), illustrated the relationships between input fMRI volumes and FC measures using both sequences (**Figure 2r**). These data suggest that SORDINO maintains sensitivity comparable to GRE-EPI BOLD-fMRI while offering additional benefits.

#### Specific Contrast Origin

Our modeling demonstrates that SORDINO is sensitive to CBV (**Figure 1e**) and tO_2_-induced T1 changes (**Figure 1f**). We conducted five experiments to empirically validate these contrasts and confirm that SORDINO does not carry conventional BOLD contrast. In **Figure 3a**, we present data from a flowing phantom inside an RF coil. The inflowing spins, owing to their initial full magnetization and the fewer RF excitations they receive before signal readout (see **Figure 1d & S2**), generate higher signals than those at steady-state, thereby elevating SORDINO signals. This characteristic sets the foundation to measure CBV changes, as increased CBV introduces more inflowing spins, enhancing imaging voxel signals due to higher vascular-space-occupancy^46,47^. In **Figure 3b**, we dissect the fraction of CBV contributions in SORDINO by comparing forepaw sensory stimulation evoked responses in rats under two coil settings: head-only excitation versus whole-body excitation. Whole-body excitation yielded approximately 40% weaker evoked responses than head-only excitation, consistent with our modeling predictions (see **Figure 1e & f and S3**). In this setting, the whole-body excitation eliminates inflow-related vascular signal enhancement, causing all blood in the subject to reach a steady-state, similar to gray matter tissue (see **Figure S3a**). As a result, any inflowing blood contribution would be minimal and negatively affect the overall SORDINO responses due to the slightly longer T1 of blood compared to tissue.

**Figure 3.**
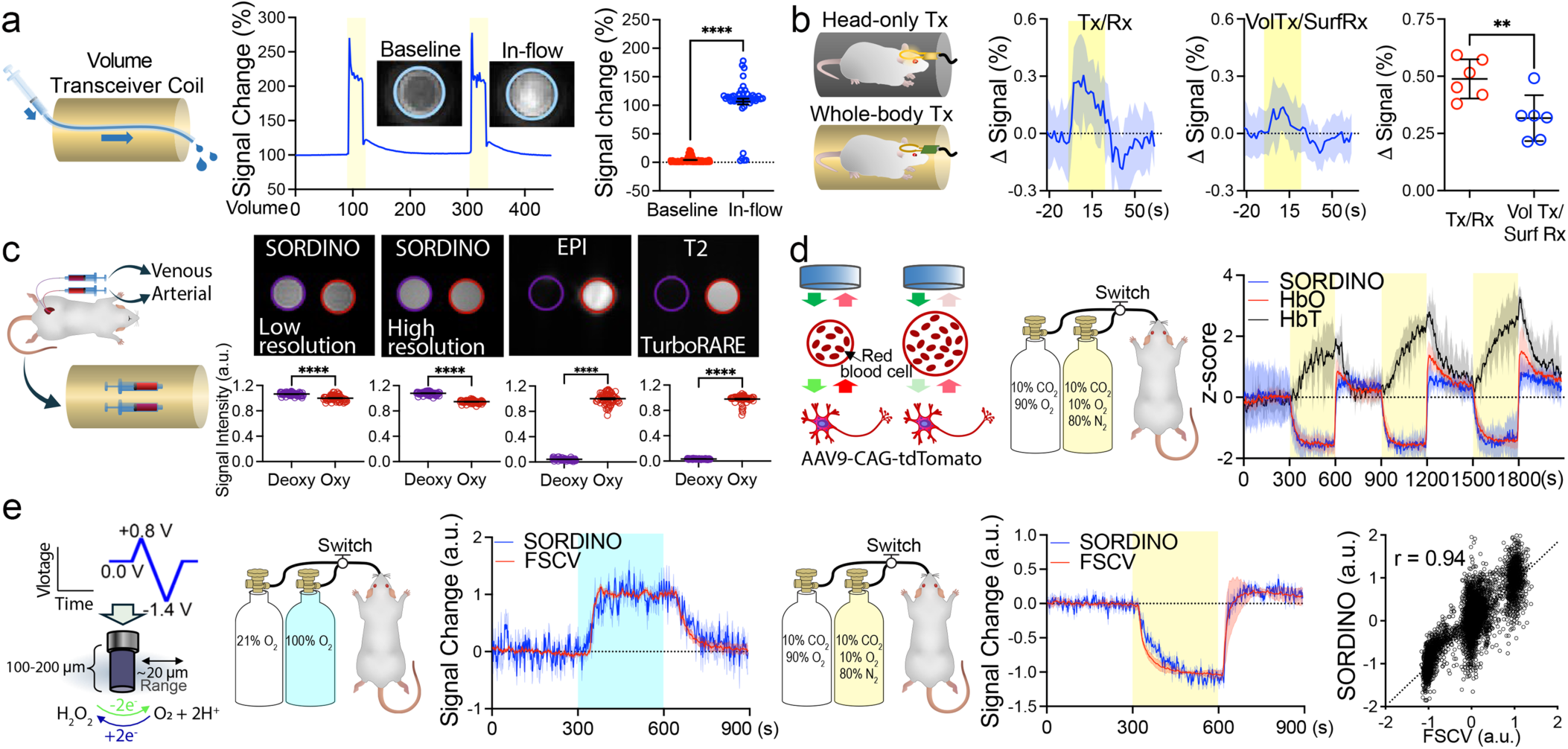
SORDINO contrast mechanisms. **(a)** SORDINO demonstrates sensitivity to inflowing spins, enabling arterial CBV contrast in vivo (n = 1 trial; 383 volumes at baseline and 52 during inflow; two-tailed unpaired t-test, t(433) = 51.08, *p* < 0.0001). **(b)** Whole-body RF excitation, compared to head-only RF excitation, eliminates CBV contributions but retains 60% of signal changes during forepaw stimulation, suggesting the presence of functional contrast mechanisms beyond CBV (n = 6 subjects; two-sided paired t-test, t(5) = 2.57, *p* = 0.049). **(c)** SORDINO lacks hemoglobin-related “BOLD” contrast. While SORDINO shows positive responses to forepaw stimulation-induced activation in S1 (see Figures 2p, **S5**, and **3b**), deoxygenated hemoglobin exhibits stronger SORDINO signals than oxygenated hemoglobin under SORDINO-fMRI parameters (left; n = 69 datapoints; two-tailed unpaired t-test, t(13) = 11.27, *p* < 0.0001) and anatomical scans with long spoke TR (right; n = 69 datapoints; two-tailed unpaired t-test, t(13) = 34.27, *p* < 0.0001). In contrast, conventional BOLD effects are reliably observed in EPI (n = 69 datapoints; two-tailed unpaired t-test, t(13) = 92.73, *p* < 0.0001) and T2-weighted scans (n = 69 datapoints; two-tailed unpaired t-test, t(13) = 126.10, *p* < 0.0001). These findings indicate that SORDINO relies on contrast mechanisms distinct from BOLD. Notably, most (∼99%) oxygen in blood is bound to hemoglobin, with minimal dissolved oxygen present. This makes the molecular oxygen effects negligible in this measurement. **(d, e)** Validation of SORDINO’s sensitivity to tissue oxygenation changes was achieved through concurrent fiber photometry and FSCV recordings. **(d)** Under hypercapnic conditions, hypoxic gas challenges resulted in increased CBV and decreased blood and tissue oxygenation, as measured by fiber photometry. The SORDINO signal closely tracked oxyhemoglobin changes rather than total hemoglobin levels, highlighting its sensitivity to non-BOLD, yet oxygen-related contrasts. Yellow boxes indicate the hypoxic gas challenge periods (n = 4, 3 unique trials per subject). **(e)** SORDINO signal changes aligned with concurrently recorded tissue oxygenation ground-truth measurements using FSCV. Left: In a hyperoxic gas challenge experiment, SORDINO signals mirrored the dynamics of tissue oxygenation (blue box indicates the hyperoxic period). Middle: During hypoxic gas challenges, SORDINO signals also closely matched tissue oxygenation changes (yellow box indicates the hypoxic period). Right: The scatter plot demonstrates a strong correlation between SORDINO signals and tissue oxygen levels (n = 3, 3 trials per animal). Results are expressed as mean ± SEM. \*\**p*<0.01 and \*\*\*\**p*<0.0001 compared with indicated groups.

In **Figure 3c**, we imaged freshly collected arterial and venous blood samples using SORDINO, GRE-EPI, and TurboRARE sequences. While the latter two sequences exhibited the typical “BOLD effects”, with higher signals in oxygenated arterial blood, SORDINO showed reversed contrast, with higher signals in deoxygenated venous blood than the oxygenated arterial blood. This contrast inversion likely arises from SORDINO’s insensitivity to paramagnetic T2* effects of deoxyhemoglobin and its efficiency in capturing the T1 shortening effects of deoxyhemoglobin^56^. These findings indicate that positive evoked responses in SORDINO (see **Figure 2p**) are not due to intravascular hemoglobin oxygenation levels, suggesting a potential solution to address the low spatial specificity and the unwanted draining vein dominance commonly seen in BOLD-fMRI^20,57–59^. In **Figure 3d**, we conducted a gas-challenge experiment with head-only RF excitation to dissociate tO_2_ from CBV. We strategically established a hypercapnic-hyperoxic baseline (10% CO_2_ and 90% O_2_), followed by a hypoxic challenge (10% CO_2_ and 10% O_2_), leading to decreased oxygenation and increased CBV. These physiological changes were validated by simultaneous measurements of SORDINO signals (head-only RF excitation), oxyhemoglobin, and total hemoglobin concentration changes (indicative of CBV) using an established fiber photometry technique^60^. The single-voxel SORDINO signals, taken at the tip of the implanted optical fiber in the cerebral cortex, closely aligned with oxygenation changes rather than total hemoglobin, featuring SORDINO’s sensitivity to non-BOLD (**Figure 3c**), yet oxygen-related contrast (**Figure 3d**). To further demonstrate the tO_2_ effects *in vivo*, in **Figure 3e**, we showcase another gas-challenge experiment, inducing positive (21% O_2_ baseline followed by 100% O_2_ hyperoxic challenge) and negative (10% CO_2_ and 90% O_2_ hyperoxic baseline followed by 10% CO_2_ and 10% O_2_ hypoxic challenge) tissue oxygen changes while concurrently acquiring SORDINO (head-only RF excitation) and FSCV-derived ground-truth tissue oxygen data^61–63^ – a setup enabled by SORDINO’s minimal gradient-induced artifacts. The single-voxel SORDINO signals at the tip of the implanted FSCV electrodes tightly coupled with tissue oxygen levels in both conditions, highlighting its robustness in detecting tissue oxygen changes. Given that oxygen is predominantly released at capillaries and arterioles^64^, in close proximity to the source of neuronal activity^19^, and that free molecular oxygen typically diffuses less than 30 μm from vascular sources^65^, SORDINO holds potential to offer improved spatial specificity over GRE-EPI BOLD-fMRI.

### SORDINO for Resting-State fMRI in Awake Mice

To demonstrate the feasibility of SORDINO for mapping functional connectivity in awake mice, we designed a dual-functional headplate coil using dielectric PCB-board material with insulated coil traces. The coil functions as the RF transceiver while also supporting the fixation of the mouse head. **Figure 4a** shows an awake mouse fitted with the coil on its head, without restraining its body or limbs. All subjects underwent a five-day habituation process, as shown in **Figure 2c**. We evaluated the effects of nuisance signal removal using a pipeline detailed in the Methods section (**Figure 4b-d**). Following pre-processing, we first performed a seed-based analysis by placing seeds in the left S1 and RSC, both of which revealed functional connectivity patterns as expected^24,27,66^. We then assessed the reliability and specificity of this SORDINO dataset in measuring functional connectivity (**Figure 4g-i**) and used independent component analysis (ICA) to extract large-scale brain networks. **Figure 4j** illustrates the representative “triple-networks”, comprising the default mode network (DMN), the lateral cortical network (LCN), and the salience network (SN) of the mouse brain^67^, which are critical in neuropsychology and related disorders^68–76^. To effectively depict large-scale functional networks and illustrate the SORDINO-derived functional connectome of the awake mouse brain, we transformed the SORDINO data into the Allen Mouse Brain Atlas space^77^. This transformation facilitates direct comparisons between SORDINO-derived functional connectivity and the structural connectivity ground truth provided by the Allen Brain Atlas (**Figure 4k**). The stress-free, distortion-free functional connectome data presented here hold promise for future comparative studies and represent a valuable resource for the neuroimaging community.

**Figure 4.**
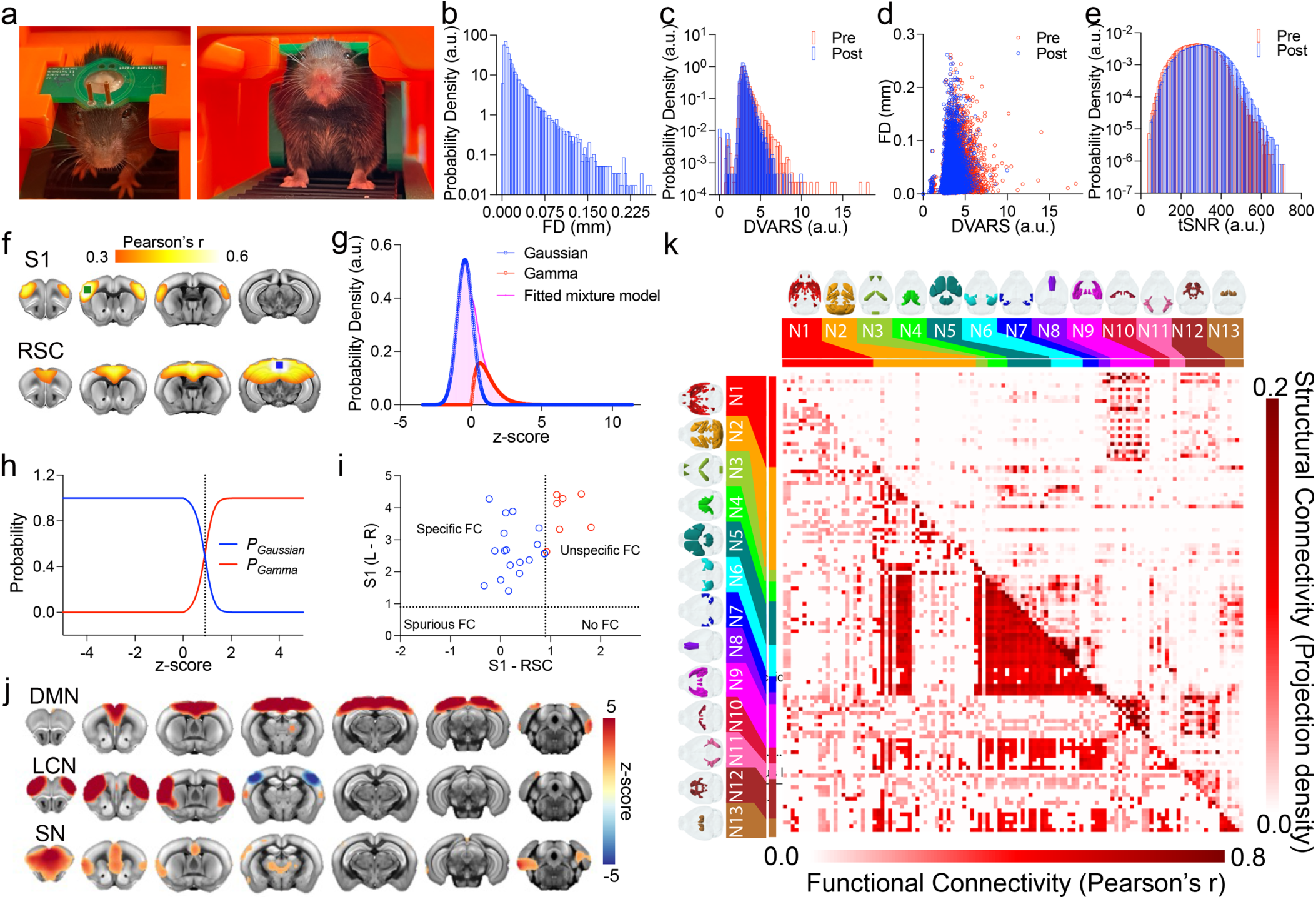
SORDINO for functional connectome mapping in awake mice. **(a)** Mouse wearing a custom headplate coil in a 3D-printed cradle for functional connectivity mapping in awake condition. **(b)** Distribution of FD data from all subject time-courses, representing FD values across every time point and displayed as a log-scale density histogram (n = 25). SORDINO data were acquired at 400 μm isotropic spatial resolution and 2 s temporal resolution, continuously for 900 volumes. The effects of nuisance removal across all subject time-courses are shown in **(c)** D-variate temporal standard deviation (DVARS) histogram, **(d)** scatterplot between FD and DVARS, and **(e)** tSNR histogram. **(f)** Voxel-wise seed-based connectivity maps from the left S1 (green) and RSC (blue) seeds across all subjects (n = 25) are shown using a second-level permutation method thresholded at *p* < 0.01 (FWE-corrected) and r > 0.3. **(g)** A Gaussian/Gamma mixture model was fitted to characterize the distribution of seed-based connectivity map. Voxels with low connectivity to the seed region were modeled by the Gaussian component (blue), while voxels with high connectivity with seed region were modeled by the Gamma component (red). **(h)** A probability curve was computed to identify the boundary between the Gaussian (blue) and Gamma (red), and the resulting boundary values were used as thresholds for determining functional specificity. **(i)** Functional connectivity specificity scatter plot showing the distribution of individual subject specificity data, defined as homotopic S1 functional connectivity across hemispheres (mirroring the seed shown in **(f)**, expected to be high in a robust dataset) relative to S1-RSC functional connectivity (expected to be low in a robust dataset). Subjects in the upper left quadrant exhibit high functional connectivity specificity, while the other three quadrants represent unspecific functional connectivity, spurious functional connectivity, and no functional connectivity. **(j)** ICA-derived “triple-network” patterns in awake mice. Coronal sections show the intrinsic large-scale networks including, DMN, LCN, SN. **(k)** Functional connectivity measured in awake mice undergoing SORDINO (n = 25) and analyzed using Allen Brain Mouse Brain ROIs is shown in the lower left of the matrix, while the Allen Atlas structural connectivity pattern is displayed in the upper right of the matrix, illustrating the functional-structural relevance of the mouse brain connectome. N1: Limbic and Associative Network, N2: Sensorimotor-Cognitive Integration Network, N3: Hippocampal-Prefrontal Associative Network, N4: Retrosplenial and Visual Network, N5: Somatosensory and Motor Network, N6: Primary Sensory Network, N7: Associative Visual Network, N8: Prefrontal Cognitive Network, N9: Limbic Emotional Network, N10: Motivational and Memory Network. N11: Visual and Entorhinal Network. N12: Subcortical Limbic Network. N13: Thalamic Relay Network.

### SORDINO for Mapping Skilled Motor Actions in Mice

Motion artifacts and stressful acoustic noise often confine conventional rodent fMRI studies to passive sensory stimulation experiments. In principle, SORDINO’s motion-insensitive (**Figure 2m-o**) and acoustically silent features (**Figure 2a-c**) provide a unique opportunity to map complex active behaviors with fMRI. To test this capability, we designed a pneumatically driven food pellet delivery system (**Figure 5a**) and trained mice to reach and grasp a food pellet using their forepaw once every 20 s under a head-fixed condition, with imaging data acquired by the same dual-functional head fixation transceiver coil plate mentioned earlier. This task requires delicate coordination of activity across the brain and triggers a motor sequence actively initiated and executed by the subjects (**Figure 5b** & **Video 4**), rather than relying on passive stimulations typically seen in most rodent fMRI studies. **Figure 5c-e** depicts the brain-wide activity changes and time-courses from various ROIs. The data revealed that system-level motor network activity consistently initiated in the contralateral motor cortex, with concurrent deactivation found in the retrosplenial cortex (RSC), a key hub of the DMN known to deactivate during task performance^71^. Subsequently, we observed robust activation in the bilateral sensory-motor cortices, dorsolateral striatum, thalamus, and cerebellum, followed by a return to baseline, reflecting behavior-related activity changes in these brain regions. Notably, the activation patterns identified with SORDINO closely align with brain regions known to play a causal role in the control of voluntary limb movements in mice^78–84^. This represents the first whole-brain map of a dexterous motor behavior in the rodent and demonstrates that SORDINO can map active behaviors, once considered extremely challenging, if not impossible, for conventional fMRI sequences.

**Figure 5.**
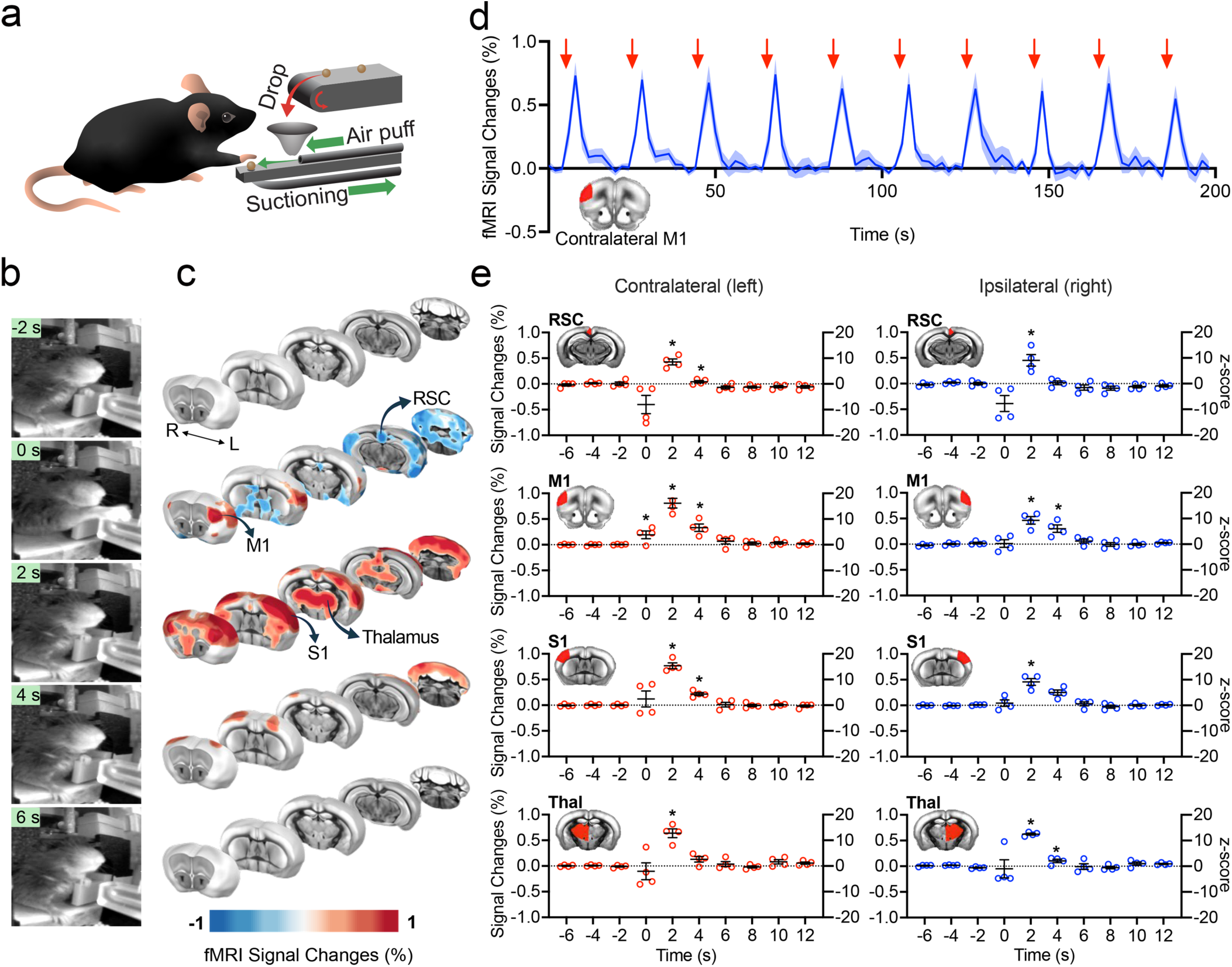
SORDINO for mapping skilled motor actions. **(a)** Custom pneumatic system enables grabbing task performance in head-fixed mice. **(b)** Mice perform reach-and-grasp tasks during SORDINO-fMRI. **(c-e)** Brain-wide activity maps and time-courses show consistent motor cortex activation and retrosplenial cortex deactivation, followed by subsequent activation of thalamus, cerebellum, and sensory cortices (n = 4, in the total of 24 fMRI sessions with 75-100 grabbing events per session). A linear mixed-effects model is utilized to compare the activity at time = 0, 2, and 4 s to the baseline (averaged activity at time = −6, −4, and −2). Results are expressed as mean ± SEM. (\**p*<0.05).

### SORDINO for Cross-Brain Coupling during Mouse Social Behavior

Leveraging SORDINO’s resistance to magnetic field inhomogeneity and motion problems, we have developed a platform that enables simultaneous imaging of two mouse brains during social behavior. This platform includes a 3D-printed cradle that mounts two awake mice face-to-face, separated by a divider plate that can be remotely withdrawn from outside the MRI scanner room (**Figure 6a** & **Video 5**). Each mouse is mounted with a head fixation plate coil, whereby RF pulses are individually transmitted and signals are received from each coil, enabling dual-brain “hyperscanning” of two socially interacting mice (**Figure 6b**). Upon removal of the divider plate, the subjects immediately display signs of social behavior, most notably in the form of increased whisking (**Video 5**). SORDINO data showed transient activations in the RSC, prelimbic (PrL), and cingulate (Cg) cortices of the DMN, and the anterior insula (AI) of the SN, peaking at 4 s; and deactivations in the M1 and the S1 forepaw (S1FL) and whisker (S1BF) regions of the LCN, peaking at 8 s (**Figure 6c**). With the divider in place, no significant inter-brain correlation was detected; however, upon its removal, mirrored inter-brain connectivity emerged in the RSC, Cg, PrL, and AI, suggesting synchronization of activity dynamics between the two brains (**Figure 6d**). Further ROI analysis revealed significant inter-brain, cross-region connectivity among M1, DMN, and SN nodes after divider removal, with no inter-brain connectivity observed during the period of having the divider in place (**Figure 6e**). Divider removal also induced a significant increase in mean cross-brain spatial correlation within the DMN and SN (**Figure 6f**). Voxel-wise analysis demonstrated consistent inter-brain synchronization within the DMN and SN regions, aligning with the partner’s RSC, Cg, PrL, AI, and M1, but not with their S1 (**Figure 6g**). This pattern suggests that inter-brain synchronization arises from perceiving the partner’s actions and broader brain states, which may be influenced by M1 activity but are not necessarily driven by M1 alone. Altogether, our results demonstrate a novel platform that opens new avenues for measuring and manipulating inter-brain communications, offering exciting possibilities for advancing neuroimaging research.

**Figure 6.**
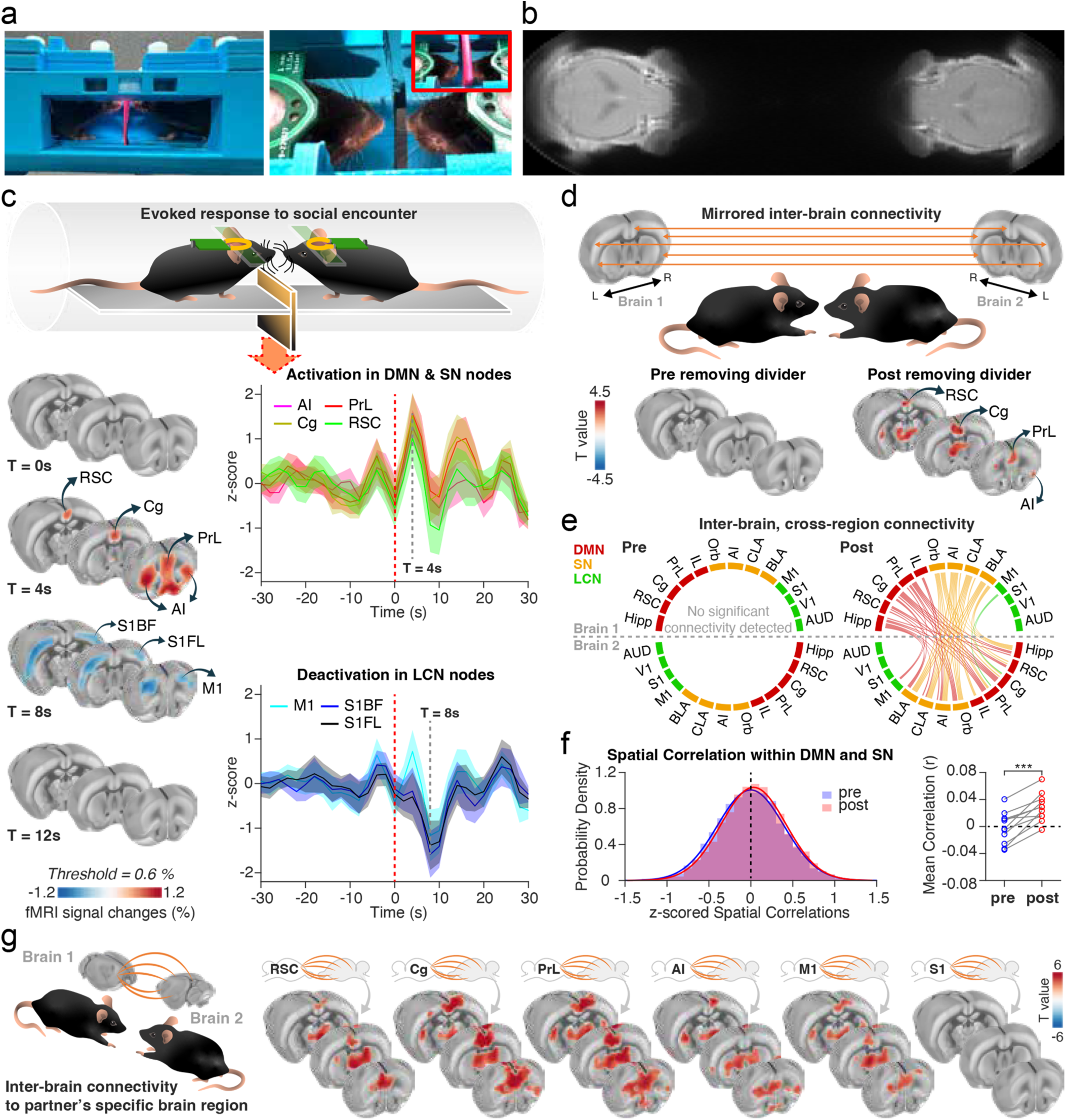
Mapping inter-brain synchronization in awake, socially interacting mice using SORDINO hyperscanning. **(a)** A 3D-printed cradle holds mice face-to-face, initially separated by a remotely retractable divider, enabling dual-brain SORDINO-fMRI hyperscanning. **(b)** A SORDINO image with an extended FOV along the gradient z-axis, showing the actual distance between the brains of two subjects. **(c)** SORDINO hyperscanning captured brain responses during social encounters initiated by divider removal (n = 22, grouped in 11 pairs, averaged signal change map, threshold = ±0.6%). Following divider removal, activations peaked at 4 s in DMN nodes (RSC, Cg, PrL) and the AI of the SN, followed by deactivations peaking at 8 s in LCN nodes (M1, S1FL, S1BF). **(d-f)** Inter-brain synchronization emerged only in socially interacting mice without the divider. **(d)** Mirrored inter-brain connectivity appeared in RSC, Cg, PrL, and AI post-divider removal (n = 11 pairs, one-sample *t*-test, *p* < .01), absent during baseline. **(e)** ROI analysis showed significant inter-brain connectivity among DMN, SN, and M1 only after divider removal, as illustrated in the right chord diagram (top and bottom hemispheres represent individual brains of paired mice, n = 22, one-sample *t*-test, *p* < 0.001, FDR corrected with Benjamini-Hochberg procedure). **(f)** Spatial correlation analysis indicated enhanced inter-brain synchrony of activation patterns within DMN and SN post-divider removal. (Left: spatial correlation distribution across pre- or post-divider-removal time period; Right: mean spatial correlation within subjects, pre- and post-divider-removal, n = 11 pairs, paired *t*-test, *p* < .001). **(g)** SORDINO signals within DMN and SN regions showed significant inter-brain synchronization with the socially interacting partner’s RSC, Cg, PrL, AI, and M1, but not with the partner’s S1 (n = 22, one-sample *t*-test, *p* < .001). AI, anterior insula; AUD, auditory cortex; BLA, basolateral amygdala; Cg, cingulate cortex; CLA, claustrum; Hipp, hippocampus; IL, infralimbic cortex; M1, motor cortex; Orb, orbitofrontal cortex; PrL, prelimbic cortex; RSC, retrosplenial cortex; S1, somatosensory cortex; V1, visual cortex.

### SORDINO for Mouse Glymphatic Dynamics

In addition to fMRI, we conducted two proof-of-concept studies to demonstrate SORDINO’s ability to robustly capture T1 changes using paramagnetic manganese ion (Mn^2+^) and gadolinium-based (Gd) contrast agents. We first modeled the contrast gain in SORDINO at 9.4T, considering a 50% of T1-shortening effect (i.e., from 1900 ms to 950 ms), which is commonly observed in manganese-enhanced MRI (MEMRI) studies^85–90^. Our modeling suggests that, with proper selection of imaging parameters, SORDINO signals are expected to be 1.8-fold stronger with manganese enhancement (**Figure S6a**). By comparing SORDINO images from the same mouse before and 7 days after subcutaneous osmotic mini-pump infusion of MnCl_2_ at a total dose of 175 mg/kg^89^, we demonstrated a robust enhancement of the SORDINO signal in key regions known to exhibit Mn^2+^ uptake. These regions include the pituitary gland, hippocampus, cerebellum, and olfactory bulb, consistent with findings in classical MEMRI studies^85–90^ (**Figure S6b, Video 6**). Next, we showcased SORDINO’s ability to rapidly image brain-wide cerebrospinal fluid (CSF) influx into the brain parenchyma using an intra-cisterna magna injection of Gd contrast (Prohance, 0.067 mM, 10 µL) – a protocol increasingly used to evaluate the glymphatic system dynamics, which are critical for brain waste clearance^91,92^. Upon Gd injection, we observed an immediate increase in SORDINO signals in the cisterna magna, followed by the fourth ventricle, third ventricle, lateral ventricle, and cortical regions (**Figure S7, Video 7**). A reversal in contrast was observed in the ventricles, where they initially appeared darker than gray matter but became brighter than gray matter following Gd injection (**Figure S7a**). These pronounced contrast changes significantly enhanced the identification of regions where Gd was distributed. In a Cartesian-sampled FLASH sequence at minimal TE (**Figure S7b**), the regions surrounding the injection site experienced signal loss due to susceptibility artifacts and/or high concentration Gd-related T2* shortening, whereas the SORDINO sequence did not show this artifact. This enabled brain-wide signal normalization to the cisterna magna following Gd injection (**Figure S7c**), thereby controlling data variability associated with injection volume variations during microinjection. Looking ahead, the silent feature of SORDINO offers the potential to image glymphatic flow without the confounding effects of anesthesia^93^. This advancement enables future investigations of the glymphatic function in both awake and naturalistic sleep conditions, thereby enhancing translational relevance to human and clinical populations known to exhibit glymphatic deficits.

## DISCUSSION

We introduce an innovative method named <u>S</u>teady-state <u>O</u>n-the-<u>R</u>amp <u>D</u>etection of <u>IN</u>duction-decay signal with <u>O</u>versampling, or SORDINO in short. Drawing from musical terminology, SORDINO signifies the muting and damping of vibrations in instruments, reflecting the features this technique provides in MRI. We evaluated SORDINO alongside GRE-EPI on various metrics, dissected SORDINO functional contrast, and demonstrated its efficacy in imaging rodent brain function in awake and behaving conditions. These studies suggest SORDINO to be a powerful technique that is silent, sensitive, specific, and resilient to artifacts commonly seen in GRE-EPI, enabling new opportunities for functional brain mapping during active behavior.

### SORDINO’s technical features

SORDINO is silent and can minimize electromagnetic interference because rapid switching of gradient polarity and gradient resetting are absent. Within an fMRI experiment, we clamp the total gradient amplitude constant, continuously rotate gradient direction, and employ a trajectory design to yield consistent and minimal gradient directional changes, never needing to reset the total gradient amplitude to zero (**Figure 1**). These features allow SORDINO to achieve an ultra-low slew rate, effectively eliminating acoustic noise and reducing subjects’ stress (**Figure 2a-c**), while also minimizing electromagnetic interference, as evidenced by electrophysiological and electrochemical recordings (**Figure 2d-f**). Currently, most simultaneous electrophysiology-fMRI studies rely heavily on post-processing and/or regression of MRI-induced noise. Our findings highlight the potential of SORDINO to facilitate simultaneous, real-time electrophysiological/electrochemical recordings during fMRI, paving the way for future close-loop studies. Additionally, the feasibility of performing cellular resolution calcium imaging during SORDINO-fMRI (**Figure 2g**) holds promises for bringing novel insights into cellular mechanisms underlying large-scale brain networks.

SORDINO is resistant to ghosting, distortion, susceptibility, and motion artifacts. Building on a 3D radial sampling strategy, SORDINO densely samples information around the k-space center, thereby distributing the energy of transient motion into thousands of encoding directions. Imaging sequences such as PROPELLER^94^ or other center-out encoding schemes^95–97^ have shown great promise in reducing ghosting and motion artifacts. As shown in **Figure 2**, SORDINO inherits these artifact-resisting properties for fMRI applications. Further, SORDINO encodes data immediately after RF excitation, leaving essentially no time for voxel dephasing. Since all datapoints are sampled using frequency encoding within a very short readout time, the phase deviation from encoding is minimized and the bandwidth is uniform along all encoded spokes. This also renders SORDINO insensitive to geometric distortion, enabling studies of brain regions susceptible to field inhomogeneity, such as amygdala in the rodent brain^66,98,99^ (**Figure 2k&l**). Consequently, this minimizes image distortion caused by field inhomogeneity in vulnerable brain regions, thereby reducing compounded errors typically encountered during rigid-body registration-based motion correction.

The most efficient sequence for mapping functional activations should minimize the time spent on contrast preparation while maximizing the time available for data acquisition. SORDINO represents a great example of this concept by acquiring data throughout the entire TR (**Figure S1a**). Specifically, it provides robust sensitivity to functional activation through several innovative strategies employed. First, imaging extremely short T2* species using a high acquisition bandwidth is not necessary for brain fMRI applications (**Figure S1h**). Therefore, SORDINO exclusively samples data on-the-ramp to afford very low bandwidth acquisition within an extremely short TR (e.g., 0.6 ms for all awake mouse studies presented in this work). Such a short TR is unattainable with conventional sequences because high acquisition bandwidth is necessary to rapidly encode the entirety of k-space between each RF pulse repetition in EPI^66^ or to accommodate multiple excitations for single spoke encoding in sequences such as MB-SWIFT^39,40^. Importantly, our data has demonstrated robust CNR compared to GRE-EPI (**Figure 2**), supporting the feasibility of achieving additional sensitivity gain with SORDINO. In addition, compared to other short-acquisition-delay sequences that rely on high acquisition bandwidth, the low bandwidth strategy affordable by SORDINO can improve SNR and reduce missing samples at the center of k-space (**Figure S1**).

SORDINO demonstrates robust sensitivity to tO_2_ or combined CBV and tO_2_ changes depending on the RF delivery strategy, setting it apart from the conventional BOLD contrast, which is heavily influenced by hemoglobin oxygenation levels^1–6^. The insensitivity to deoxyhemoglobin’s paramagnetic effects, together with the high spatial specificity of tO_2_^19,64^ and CBV changes^58,100^, suggests that SORDINO can offer more precise localization of neuronal activity by minimizing the influence of draining veins, a common issue in BOLD-fMRI^6,19^. By testing head-only and whole-body RF coil excitation settings within the same scanning session, we were able to examine the CBV contribution to SORDINO signals under head-only RF excitation. The experiments demonstrated that head-only RF excitation can approximately double the fMRI response amplitude by capitalizing on the inflow arterial signal enhancement. While this is beneficial for improving data acquisition efficiency, this extra sensitivity gain may be dependent on the blood transit times. For instance, it has been shown that the frontal brain regions generally exhibit a transit time delay of approximately 0.04 s compared to posterior brain regions^101^. This highlights the need for future studies to thoroughly investigate how region-dependent blood transit times may influence SORDINO sensitivity. In both simultaneous photometry-fMRI and FSCV-fMRI experiments, we observed SORDINO’s sensitivity to oxygenation, which aligns well with the modeled T1 shortening effects of molecular oxygen. Given that oxygen is predominantly released at capillaries and arterioles near neuronal activity sites^19,64^, SORDINO’s responses are likely to originate from regions closer to the sources of neuronal activity than BOLD-fMRI. It should be noted that while SORDINO offers significant advantages, its application in very high temporospatial resolution laminar fMRI studies remains challenging. This is primarily due to the use of nonselective RF pulses and SORDINO’s 3D imaging nature, which would require impractically long imaging times if resolution were substantially increased. While undersampling can help mitigate prolonged imaging times, a high undersampling factor introduces artifacts (**Figure S8**). In this work, we focus on demonstrating the feasibility of SORDINO and providing a fair comparison against EPI, while leaving the exploration of acceleration options and regularized reconstruction approaches for future studies. Translating SORDINO to human MRI systems where multi-channel coils enable additional acceleration options, could be critical for further demonstrating its capabilities and spatial specificity.

Given that tissue T1 increases at higher magnetic field strengths while the molecular oxygen relaxivity remains relatively unchanged^102^, we expect SORDINO to perform better at higher magnetic fields (**Figure S9**). This enhanced sensitivity to T1-shortening effects, alone with the known SNR benefits from increased polarization of spins and reduced thermal noise^103^, highlights the potential utility of SORDINO at high magnetic fields. fMRI applications at high field strengths often face challenges such as susceptibility artifacts, eddy currents, and difficulties encoding shortened T2* signals in time – problems to which SORDINO is relatively immune.

### SORDINO applications

SORDINO represents a transformative tool for conducting fMRI studies in awake mice, offering unprecedented opportunities to map brain function during resting-state, naturalistic behaviors, social interactions, and glymphatic dynamics, previously constrained by the limitations of conventional MR methods. In a series of four proof-of-concept studies, we demonstrated that SORDINO enabled FC mapping in awake mice with significantly reduced habituation time from the month-long process typically required^27–29^ to just 5 days. We attribute this improvement from SORDINO’s elimination of the acoustic noise and gradient vibration inherent in conventional EPI-fMRI, which necessitate extended habituation to mitigate stress in the animals. The shortened habituation period aligns with what is observed in head-fixed optical imaging studies, where the animals do not encounter noisy environments and have the ability to move^104–107^. SORDINO’s streamlined FC mapping significantly enhances the accessibility of awake rodent fMRI to the broader research community. This development is particularly impactful given that much of current rodent fMRI is still performed under anesthesia^24^ – a limitation that the field is keen to overcome.

We demonstrated that SORDINO effectively images brain activity during a forepaw-reaching task in mice using a custom-designed, MR-compatible food pellet delivery system. This capability to capture active, voluntary behavior overcomes traditional challenges of fMRI in moving subjects, enabling naturalistic fMRI studies of motor behavior-driven brain activity changes. SORDINO’s insensitivity to electromagnetic interference further allows for seamless integration of complementary modalities, facilitating circuit-based behavior augmentation or piloting novel therapies for movement and mental disorders while establishing their causality to brain networks. Additionally, we extended SORDINO’s application to simultaneous imaging of two mouse brains during social behavior, revealing cross-brain coupling in large-scale brain networks. This platform paves the way for novel research opportunities, such as using closed-loop setups to record and feed signals between subjects while simultaneously imaging brain-wide activity. The ability to study vocalization and communication between subjects without the confounding effects of MRI-related acoustic noise also offers exciting potential for advancing knowledge of these sophisticated behaviors. Further, we showcased SORDINO’s versatility beyond traditional fMRI applications by mapping the glymphatic system, which is gaining significant attention for its role in neurodevelopmental and neurodegenerative disorders. Using Gd contrast, SORDINO provided clear, artifact-free imaging of CSF influx, distinct from traditional methods that suffer from susceptibility artifacts and Gd-related T2 shortening. The silent imaging features of SORDINO are particularly advantageous for studying glymphatic flow in awake, sleep, and naturalistic conditions, a significant step forward in understanding glymphatic system’s role in brain health without confounds associated with anesthesia.

## CONCLUSION

We describe the theoretical framework and implementation of SORDINO, identifying robust in vivo parameters and benchmarking them against modeled biophysical properties and various ground-truth validations. This work empirically demonstrates SORDINO’s sensitivity to tO_2_ and CBV, suggesting its spatial specificity, and highlights several novel applications enabled by this technique. The information, raw data, and sequences provided in this manuscript are intended to facilitate future use of SORDINO within the fMRI community and advance the exploration of functionally and behaviorally relevant brain circuits in awake animal subjects. This study pushes the boundaries of fMRI by offering a silent, sensitive, and specific method with reduced artifact interference. The insights gained here are anticipated to have broader implications beyond fMRI, with these benefits translating to dynamic contrast-enhanced MRI^108,109^ and molecular MRI^110^ studies in awake, behaving animal subjects.

## METHODS

Male C57BL/6J mice (n = 147; 23-32 g), male Sprague-Dawley rats (n = 43; 232-500 g) were acquired from Charles River (Wilmington, MA, USA). Animals were given food and water ad libitum and kept on a 12-hour light/dark cycle. In addition to MRI, we integrated established neuroscience tools for recordings during SORDINO-fMRI. These tools included electrophysiology, FSCV, miniscope calcium imaging, and spectral-fiber photometry. Stock AAV vectors were purchased directly from Addgene. Given the variety of approaches used in this manuscript, we structured the methods into four main categories: (1) MRI hardware, sequence, and modeling, (2) Phantom experiments and SORDINO sequence characteristics, (3) Rat experiments, and (4) Mouse experiments. Methods within each category follow the order in which they are presented in the main text. Python (version 3.7) and the NumPy^111^, MATLAB (R2024a) and GraphPad Prism 10 were used for data analysis.

### MRI Hardware, Sequence, and Modeling

All MRI experiments were performed on a Bruker 9.4 T/30 – cm horizontal bore system, with a BFG 240/120 gradient insert (Resonance Research Inc, Billerica, MA) and an AVNEO console, configured with a maximum gradient amplitude of 922 mT/m.

#### SORDINO sequence

The SORDINO sequence demonstrated in this manuscript was programmed on Bruker ParaVision 6 and 360, building upon the default Bruker ZTE sequence. Software and hardware configuration remained unchanged between the two versions, ensuring consistent sequence performance throughout this work. The general framework of the single spoke acquisition in SORDINO is shown in **Figure 1a**. Following a one-time initialization of the encoding gradient, a short, hard RF-pulse is applied. This RF-pulse, lasting only a few µs, enables a short-acquisition delay.

#### Clamping the encoding gradient slew rate

The primary aim of SORDINO was to develop a sequence with minimal gradient switching to minimize acoustic noise and electromagnetic interference. In MRI, fast switching currents induce Lorentz forces in the gradient coils, causing vibrations in the surrounding structures and resulting in the loud knocking sound characteristic of EPI. The rapidly varying magnetic fields induced by gradient switching generate concomitant currents in metallic components such as wires and electrode leads, making electrical recordings challenging without proper artifact regression or interleaving the MRI acquisition. Conventional ZTE realizes low gradient switching by ramping the encoding gradient before RF-application while keeping the gradient switched on throughout the acquisition, with only incremental adjustments in amplitude between spokes. By default, the ZTE gradient amplitude is adjusted over the configured ramp time, which was 0.21 ms on our system. SORDINO achieves a significantly lower slew rate than ZTE by nearly clamping the gradient slew rate constant throughout acquisition. This was accomplished by adjusting where the program initiates gradient ramping and extending ramp time to its maximum possible duration. However, uninterrupted ramping was not feasible on our system due to potential memory constraints with the sequencer. Instead, a 10 µs plateau was introduced between spokes before the gradient resumed ramping. The low slew rate strategy of SORDINO offers an additional potential benefit for fMRI. Enabling continuous ramping throughout acquisition eliminates delays associated with switching the gradient ramping on and off, allowing for a lower acquisition bandwidth. A reduced bandwidth proportionally decreases noise variance in the sampled data and increases SNR by reducing the thermal noise component of fMRI data^112^. Furthermore, since gradient strength decreases proportionally with receiver bandwidth, a lower bandwidth enables sampling closer to the center of k-space. This is because fixed delays in the sequence – such as RF-pulse duration, ADC initialization and coil ringdown – are independent of bandwidth.

#### RF-spoiling and pulse length

Because our goal was to minimize gradient ramping and maximize acquisition duration, we did not employ gradient-spoiling in SORDINO. Instead, we used RF-spoiling wherein the phase of the RF-pulse was incremented by 117°, upon each repetition of the excitation pulse, which mixes transverse coherence pathways, reducing the formation of spurious echoes and the temporal variability of the signal^113^. In this manuscript, spurious echoes refer to unintended echo formation regardless of the mechanism. Throughout all experiments, we selected a 4 µs RF-pulse length as a compromise between minimizing the acquisition delay, ensuring RF power stability, and maintaining sufficient B_1_ homogeneity.

#### Spoke-oversampling

On our system, the first sample in all SORDINO acquisitions was corrupted due to a digital filter group delay. With spoke-oversampling, each sample represents a smaller time step, meaning that the number of delayed samples corresponds to a shorter duration. This enables the acquisition of samples closer to the center of k-space, mitigating the impact of group delay on image quality. Since the reconstructed FOV is inversely proportional to sampling distance, oversampling provides the flexibility to reconstruct images that mitigate aliasing from unwanted signals outside the nominally prescribed FOV. This is particularly important for SORDINO, which is sensitive to signals from short-T2 materials, such as those from coil insulation materials (**Figure S1h**). All experiments described in this manuscript used an oversampling factor of 8 as a compromise between data size and mitigating the effect of digital filter group delay.

#### Comparison of gradient slew rates in SORDINO, ZTE and EPI acquisitions

The slew rates of EPI, conventional ZTE, and SORDINO were compared based on the acquisition parameters used in rats, as described in *Comparing Sensitivity of SORDINO and EPI*, considering gradient increments in a single direction. The relationship between the change in k-space position, gradient strength and ramp time is expressed by: Δ*k*_*r̂*_= *γG*Δ*t*, where Δ*k*_*r̂*_(m^-1^) is the change in k-space position, representing movement along a spatial frequency range; *γ* (MHz/T) is the gyromagnetic ratio; *G* (T/m) is the gradient amplitude; and Δ*t* is the gradient ramp time. Rearranging this equation to determine the gradient amplitude required to traverse a specified distance in k-space and normalizing the result to units per second yields the slew rate: 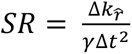. Under our standard rat SORDINO imaging parameters, with a TR of 1.82 ms, a temporal resolution of 2 s, and an isotropic spatial resolution of 0.6 mm^3^, the k-space extent is 1,666.67 m^-1^. As SORDINO encoding progresses from the center of k-space to the periphery, the signal is encoded over 833.34 m^-1^ per projection, with a ramp time of 1.81 ms. Although the total gradient slew rate and gradient amplitude are clamped across the acquisition of a volume, they are not identical for any given encoding direction. The mean incrementation between each projection is 0.00038 T/m, corresponding to a slew rate of 0.35 T/m/s. The conventional ZTE sequence on a Bruker system differs from SORDINO in that the encoding gradients are incremented between successive spokes. With a calibrated ramp time of 0.21 ms, the mean gradient incrementation of an equivalent ZTE sequence is 0.0033 T/m, yielding slew rates of 15.66 T/m/s — two orders of magnitude higher than SORDINO. In our rat fMRI experiments using EPI, a 2D acquisition with an in-plane spatial resolution of 0.6 mm also has a k-space extent of 1,666.67 m^-1^. The acquisition used a matrix size of 42 and employed a 50 µs gradient blip between successive frequency encoding steps, resulting in a phase encoding gradient amplitude of 0.018 T/m and a slew rate of 355.84 T/m/s. With a 250 kHz receiver bandwidth, sampling 44 points along the frequency encoding direction takes 0.176 ms. This corresponds to a gradient amplitude of 0.22 T/m and a slew rate of 1263.62 T/m/s — five and three orders of magnitude higher than SORDINO and ZTE, respectively.

#### Bloch equation modeling of SORDINO signal

To understand the characteristics and potential sources of the SORDINO functional contrast, we carried out Bloch equation modeling. The Bloch equations describe how the SORDINO signal evolves over time in response to elements of the pulse sequence and relaxation^114^. We analyzed the Bloch equations in the rotating frame and did not consider the effect of B_1_ or gradients on precession. Further, our simulations focused solely on isochromats at isocenter, disregarding off-resonance precession. With these assumptions, the evolution of the net magnetization vector, **M**, as a function of time is: **M**(t) = **AM** + **B**

Where:

- **A** is a 3×3 matrix representing relaxation: 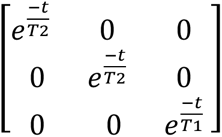,
- **B** is a 3×1 vector representing recovery of the longitudinal component of the net magnetization vector towards equilibrium: 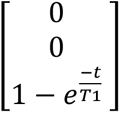.

RF-excitation was modeled as a rotation about the y-axis: 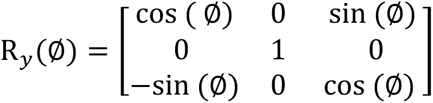

Where:

- ∅ is the FA.

Since the acquisition delay of SORDINO – on the order of microseconds – is negligible compared to the T2* of the rodent brain, the SORDINO signal was taken as the transverse component of **M** immediately following excitation. Consequently, precession was only considered over the repetition time (TR). In some cases, as RF-spoiling was implemented (**Figure S1b**), which is expected to significantly reduce formation of spurious echoes, perfect spoiling was assumed and the transverse component of **M** was set to 0 at the end of every TR interval. Putting these components together, the **M** after n repetitions can be expressed as: 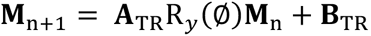.

#### Modeling of SORDINO functional contrast

Bloch equation modeling was carried out to estimate the source of the SORDINO functional contrast, considering changes in inflow-enhanced CBV and tissue oxygen levels (**Figure 1e-f**). Since RF-spoiling was implemented (**Figure S1b**)^113^, which is expected to significantly reduce formation of spurious echoes, perfect spoiling was assumed. The voxel signal was estimated by modeling the extravascular and intravascular compartments separately. Assuming the voxel-level T1 of 1900 ms, based on our own measurements (data not shown), a blood T1 of 2429 ms at 9.4 T^115^ and a vascular volume fraction of 5%, we estimated that the T1-value of extravascular tissue is 1878 ms. With head-only excitation, the inflow effect leads to apparent T1-shortening of the intravascular signal, resulting in a stronger signal than the stationary extravascular tissue at steady-state as flowing spins are subjected to RF-pulses only when within the sensitivity region of the transmitter. If the arterial transit time represents the duration flowing blood remains within the sensitivity region of the transmitter before reaching the brain, the blood signal will be in a dynamic state influenced by the arterial transit time, TR and flip angle of the sequence. Here, we assumed an arterial transit time of 280 ms^116^, a TR of 1 ms and flip angle of 3°. We estimated that the voxel signal at baseline comprises 95% extravascular tissue signal at a steady-state and 5% intravascular signal in a dynamic state. To model the effect of inflow-enhanced CBV, we assumed a local blood flow acceleration of 1.5 mm/s over an activation radius of 1 mm from a baseline flow of 10 mm/s^117^. This results in regionally accelerated blood being subjected to 13 fewer RF-pulses, along with a 20% increase in CBV. We estimated the effect of alterations in tO_2_ levels, guided by invasive measurements^48^ and molecular oxygen spin-lattice relaxivity studies^49^. Our model restricted the influence of tO_2_ to the extravascular compartment as physiological changes in blood oxygenation are not expected to influence T1 significantly^56^. Given that the change in T1-relaxation rate is linearly dependent on concentration and assuming a 30 µM activation-induced increase in pO_2_ and a r1 of 0.3 mM^-1^s^-149^, we estimated that the T1 of the extravascular tissue would be reduced by 37 ms. Together, these effects are expected result in an increase in the SORDINO signal. To model the effect of whole-body excitation, the hardware setup most commonly used in the clinical environment, we neglected the effect of inflow on blood signal in SORDINO contrast. Under this condition, we expect that an increase in CBV would result in a small decrease in the SORDINO signal due to the shorter T1 of blood compared to tissue. Modeling suggests that the combined effect of CBV and tO_2_ on the SORDINO signal would result in an increase in signal with magnitude approximately 50% of that observed under head-only excitation (**Figure S3**).

#### TR and flip angle selections in SORDINO

To understand how the physical properties of the brain and imaging parameters influence the temporal evolution of the SORDINO signal, we modeled the SORDINO signal from the first excitation to the steady-state, assuming perfect spoiling (**Figure S4a-c**). This was done across five TRs (0.5, 0.7, 1, 2 and 3 ms), five flip angles (1°, 2°, 4°, 6° and 8°) and five baseline T1 values (500, 1000, 1500, 2000, and 3000 ms), while keeping non-varying parameters constant (TR = 2 ms, flip angle = 2° and T1 = 3000 ms). Our findings indicated that the steady-state SORDINO signal is sensitive to both TR and flip angle. Next, we sought to understand how SORDINO contrast varies with these imaging parameters (**Figure S4e-f)**. Although a longer TR increases steady-state signal magnitude, a shorter TR allows for more spokes to be acquired in a given time. This reduces noise variance and is expected to minimize undersampling artifacts in SORDINO images. To account for this effect, we modeled contrast efficiency, defined as the raw contrast at steady-state divided by 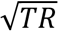^53^ at five TRs (0.5, 0.6, 0.7, 0.8, 0.9 and 1 ms) and five flip angles (1°, 2°, 3°, 4°, 5° and 6°). We assumed a 2% reduction in T1 from a physiological baseline of 1900 ms, with fixed parameters of TR = 0.7 ms and flip angle = 4°. Together, these models revealed that the sensitivity of SORDINO to T1 changes is dependent on both TR and flip angle. To identify an optimal parameter range for fMRI, we modeled the SORDINO efficiency under the assumption of perfect spoiling, across ten equally spaced TRs between 0.6 and 2.4 ms and six flip angles between 1 and 6° (**Figure S4g**). The model assumed a 2% decrease in T1 from a physiological baseline of 1900 ms, based on our own measurements (data not shown).

#### SORDINO efficiency with MEMRI

To guide imaging parameter selection for proof-of-concept MEMRI experiments, we modeled SORDINO efficiency as described in *TR and Flip Angle Selections in SORDINO* at 9.4 T (**Figure S6a**). Based on literature findings, we assumed a 50% reduction in T1^89^ from a baseline of 1900 ms. SORDINO efficiency was calculated across eight equally spaced TRs and flip angles ranging from 0.7 to 1.4 ms and 1 to 8°, respectively.

#### SORDINO-fMRI efficiency as a function of field strength

To estimate the performance of SORDINO-fMRI at various magnetic field strengths, we calculated efficiency as described in *TR and Flip Angle Selections in SORDINO* (**Figure S9).** In all cases, we assumed a 30 µM^48^ activation-induced increase in tO_2_ and a r1 of 0.3 mM^-1^s^-149^ but different baseline T1 values. Guided by the literature, we assumed T1 values of 1035, 1272, 1600, 1900 and 2030 ms at field strengths of 1.5^118^, 3^119^, 4.7^120^, 9.4 and 17.6 T^121^, respectively.

### Phantom Experiments and SORDINO Sequence Characteristics

To generate hypotheses about SORDINO’s ability to detect functional activations and its potential underlying contrast mechanisms, we conducted experiments to assess the temporal properties of the SORDINO signal and evaluate its suitability for functional imaging.

#### Assessment of gradient ramp time on SORDINO signal

In preliminary experiments, we assessed whether the strategy of increasing gradient ramp time (i.e. reducing slew rate) impacts SORDINO image quality (**Figure S1c**). Using a homemade 1 cm ID surface transceiver, we acquired SORDINO images of deionized water and CuSO_4_ phantoms at a 3 s temporal resolution with the following parameters: TR = 1ms, flip angle =3°, receiver bandwidth = 99 kHz, and spokes = 3042. Data were acquired at 9 equally spaced ramp times between 0.1 and 0.9 ms, as well as an additional acquisition at 0.99 ms. Six repeated measurements of the phantoms were acquired at each ramp time. Image quality was assessed by computing tSNR and CNR metrics.

#### Assessment of receiver bandwidth on SORDINO tSNR

After determining that extending the gradient ramping time does not adversely impact data quality, we assessed whether the low acquisition bandwidth of SORDINO increases tSNR as expected. We compared SORDINO acquisitions at various receiver bandwidths in deionized water phantoms (**Figure S1d**) and a perfusion-fixed mouse brain **(Figure S1i**). SORDINO images of deionized water were acquired using a homemade 1 cm ID surface transceiver at a 3 s temporal resolution with the following parameters: TR = 1ms, flip angle = 3°, pulse length = 4 µs, spokes = 3042, FOV = 40 mm^3^, matrix size = 60^3^, and 80 repetitions at steady-state at two receiver bandwidths – 32 and 100 kHz. Six repeated measurements of the phantoms at each receiver bandwidth were acquired. The differences between each pair of measurements were tested for normality using a Shapiro-Wilk test, and the difference between the two measurements was assessed with a paired sample t-test. Next, we acquired SORDINO images of a perfusion-fixed mouse brain at receiver bandwidths of 32, 50, 75 and 100 kHz. The common imaging parameters were: TR = 2.5 ms, flip angle = 10°, spokes = 1966, FOV = 40 mm^3^, matrix size = 60^3^ and 300 repetitions at steady-state. Six repeated measurements were taken at receiver bandwidths between 32 and 75 kHz, and five at 100 kHz.

#### Assessment of RF spoiling on FID signal using single-spoke acquisition

RF-spoiling minimizes the unintended magnetization coherence formation, resulting in the generation of echoes, hereafter referred to as spurious echoes. We qualitatively assessed the effect of RF-spoiling on echo formation by implementing a single-spoke acquisition. This model comprises periodic repetitions of identical RF-pulses and gradients, leading to the formation of stimulated echoes at predictable intervals. We chose this model because it reliably produces strong and predictable stimulated echoes, allowing us to assess the maximum effect of RF-spoiling. The single-spoke acquisition was implemented by maintaining constant encoding gradient amplitudes. RF-spoiling was implemented by incrementing the phase of the RF-pulse by 117° upon each subsequent excitation. The acquisition parameters were as follows: TR = 0.517 ms, flip angle = 4°, 60 samples per projection, 120760 repetitions and receiver bandwidth = 100 kHz, x and y-gradient amplitudes = 1.92 × 10^-4^ T/m and z-gradient amplitude = 0.013 T/m. The FID amplitude was defined as the absolute value of the first sample of each projection acquired, following the dead time gap.

#### Analysis of water phantom data

Data from the three phantom experiments described above were reconstructed using an in-house script with the Berkeley Advanced Reconstruction Toolbox (BART)^122^. Python (version 3.7) and the NumPy^111^, Statsmodels^123^, SciPy^124^ and Pingouin^125^ toolboxes were used for the analysis of phantom data. ROIs were manually drawn using ITK-SNAP^126^. The effect of ramp time on tSNR and CNR (**Figure S1c**) was assessed using a repeated-measures ANOVA. Mauchly’s test confirmed that the variances of differences between all groups were equal. Tukey’s HSD was performed for post-hoc analysis. tSNR was calculated at the voxel-level by dividing the mean signal intensity in each phantom across time by the standard deviation of the signal. CNR was calculated by subtracting the mean signal intensity of the water phantom across time from the corresponding CuSO_4_ signal intensity and dividing by the standard deviation of the signal across time. The effect of receiver bandwidth on deionized water phantom tSNR was assessed using a two-sided paired t-test (**Figure S1d**). The differences between each pair of measurements were tested for normality using a Shapiro-Wilk test. A linear mixed-effects model was used to assess the effect of receiver bandwidth on tSNR **(Figure S1i)**. Receiver bandwidth was modeled as a fixed effect and trial number was modeled as a nested random effect to account for repeated measurements. Tukey’s HSD was performed for post-hoc analysis.

#### Assessment of acoustic noise

Acoustic Noise measurements were taken for the scanner idle-state, SORDINO (volume TR = 2000 ms, spoke-TR = 0.625 ms, spokes = 3200, receiver bandwidth = 62.5 kHz, flip angle = 3°, matrix size = 64^3^, FOV = 25.6 mm^3^, voxel size = 400 µm^3^), ZTE (volume TR = 2028 ms, spoke-TR = 1.54 ms, spokes = 1320, receiver bandwidth = 62.5 kHz, flip angle = 3°, matrix size = 64^3^, FOV = 25.6 mm^3^, voxel size = 400 µm^3^), and single-shot EPI (volume TR = 2000 ms, TE = 14 ms, bandwidth = 250 kHz, flip angle = 70°, matrix size = 64 x 64 x 22, FOV = 25.6 x 25.6 x 8.8 mm, voxel size = 400 µm^3^) sequences using a Lavalier microphone (Lavalier, NY), and Audacity software (Muse Group, Limassol, Cyprus) at a sampling rate of 30 kHz (**Figure 2a**). The microphone was taped to a custom-built MRI compatible animal cradle and placed at the magnet isocenter. The recorded acoustic noise was calibrated using a frequency response curve with known volume monotones at 100 dB across frequencies of 0.2, 0.5, 1, 1.5, 1.75, 2, 3, 4, 6, 8, 16, 18 and 23 kHz. Morlet Wavelet Transformation was applied to convert the raw data into a time-frequency representation. For statistical analysis, data were converted to ΔdB, which was calculated by subtracting the scanner idle state sound pressure level from the SORDINO, ZTE and EPI recordings. The effect of sequence on sound pressure level was tested using a one-way ANOVA, followed by Tukey’s HSD test for post-hoc comparisons.

#### Assessment of electromagnetic interference using electrophysiology

Electrophysiological spiking activity signals were generated using a Blackrock Digital Neural Signal simulator (Blackrock Microsystems, Salt Lake City, UT). The simulator was connected to an MRI-compatible headstage that allows for flexible switch between two modalities: electrophysiological and electrochemical recordings (University of North Carolina CRITCL Facility, Chapel Hill, NC). The wire-junction between the headstage and simulator was attached to a water phantom placed inside the MRI bore using a 3D-printed cradle. The electrophysiological signals were recorded using a 16-channel Blackrock Cerebus System (Blackrock Microsystems, Salt Lake City, UT). Spiking activity data were recorded at a sampling rate of 30 kHz, with bandpass filtering between 0.25 and 5 kHz, during 1-minute periods, with each sequence initiated 30 s after the start of data acquisition. Electrophysiology data were preprocessed using the NeuroExplorer 5 data analysis software package (Nex Technologies, Colorado Springs, CO). Briefly, 1-minute voltage versus time courses were plotted to identify the effects of each sequence application. Simulated spikes were sorted and averaged during baseline recording (30 s before sequence initiation) to verify the signal of interest under high magnetic field condition. Similarly, spikes were sorted and averaged during control and SORDINO conditions.

#### Assessment of electromagnetic interference using FSCV dummy cell

A “dummy cell” was fabricated to characterize FSCV signals recorded within the MRI bore while three sequences were applied. The cell consisted of a 30 kΩ resistor and a 0.33 nF capacitor connected in series on a circuit board^127^, mimicking solution resistance and the double-layer capacitance of a carbon fiber electrode, respectively. During the experiment, the cell was attached to a water phantom, connected to the MRI-compatible FSCV headstage and placed inside the bore. Dummy cell-based FSCV signals were collected in 30 s recordings per sequence, including periods before and during the sequence application.

#### Assessment of ghosting artifacts in water phantom

Ghosting was measured following the guidelines of the American College of Radiology^128^ (**Figure 2i**). Nineteen volumes of a deionized water phantom were acquired using a 35 mm volume coil (Bruker Corp., Billerica, MA, USA). The imaging parameters for SORDINO were: TR = 1 ms, FA = 3°, receiver bandwidth = 300 kHz, spokes = 2000, matrix size = 100^3^, and FOV = 60 mm^3^. The imaging parameters for EPI were: TR = 2s, TE = 14 ms, flip angle = 70°, receiver bandwidth = 300 kHz, matrix size = 100^3^, FOV = 60 mm^3^, slices = 43 and slice thickness = 0.6 mm. Percent signal ghosting was quantified as: 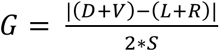, where *S*, is the signal intensity of the phantom, and *D*, *V*, *L* and *R* are the background noise signal intensities at the dorsal, ventral, left and right sides of the phantom, respectively. The differences between each pair of measurements were tested for normality using a Shapiro-Wilk test, and the difference in ghosting between the two SORDINO and EPI was assessed using a paired sample t-test.

#### Assessment of distortion in phantom

We quantified the sensitivity of SORDINO and Spin-Echo EPI sequences to geometric distortion by comparing them to Cartesian sampled spin-echo sequence (**Figure 2j**), which serves as a reliable ground truth. A spin-echo (TurboRARE), sequence is a reliable reference due to its well-documented insensitivity to static magnetic field inhomogeneities. A spin-echo EPI sequence was compared to SORDINO, as GRE-EPI acquisitions suffered from severe distortion that precluded analysis. Images of a resolution phantom were acquired using a 72 mm volume coil (Bruker Corp., Billerica, MA, USA). SORDINO parameters were: TR = 0.625 ms, FA = 3°, spokes = 31756, receiver bandwidth = 125 kHz, matrix size = 100^3^ and FOV = 40 mm^3^. TurboRARE parameters were: TR = 3600 ms, TE = 31.5 ms, echo spacing = 10.5 ms, RARE Factor = 8, matrix size = 100^2^, FOV = 40 mm^2^, slices = 32 and slice thickness = 0.4 mm and 32 averages. Spin-Echo EPI parameters: TR = 2000 ms, TE = 14 ms, receiver bandwidth = 259 kHz, matrix size = 100^2^, FOV = 40 mm^2^, slices = 32 and slice thickness = 0.4 mm. For each structure in the resolution phantom, circular features were detected using the Hough transform-based algorithm in T2-weighted TurboRARE, EPI, and SORDINO images. Detected circles were manually inspected, and outliers were removed to ensure accurate localization. To assess spatial consistency across imaging modalities, the largest center circle in each modality was designated as the reference center. The Euclidean distances from this central circle to all other detected circles within each modality were computed. The T2-weighted TurboRARE sequence served as the standard reference, and the differences in distances between T2 and the other modalities (SORDINO and EPI) were calculated. Specifically, for each corresponding circle, the difference between its distance from the center in SORDINO/EPI and its distance in T2 was determined. These distance deviations quantified spatial distortions between imaging modalities. The differences in spatial localization (i.e., distortion) between the SORDINO and Spin-Echo EPI sequences were assessed using a two-tailed two-sample *t*-test.

#### Inflow phantom

The sensitivity of SORDINO to inflow was assessed using a flowing phantom comprising PE-50 tubing connected at one end to a syringe and filled with deionized water (**Figure 3a**). The tubing was placed inside a 35 mm volume transceiver, and SORDINO images were acquired using the default mouse fMRI parameters described below. The experiment followed a block design comprising 450 volumes, with two one-minute inflow blocks starting at the 90^th^ and 300^th^ volumes. During analysis, signals were extracted from 383 baseline volumes and 52 inflow volumes, focusing on the central view of the 3D image at the magnet isocenter. Individual time series were converted to percent signal change by subtracting the baseline signal intensity from the time course, followed by a division of the mean baseline signal intensity and multiplication by 100. All data were then normalized to the mean baseline signal intensity across all trials. The effect of inflow on the SORDINO signal was assessed using a two-tailed unpaired t-test.

#### Freshly collected blood phantoms

We evaluated the sensitivity of SORDINO, EPI and RARE sequences to the BOLD contrast by imaging freshly collected arterial and venous blood samples from a single rat, leveraging the natural difference in oxygenation between venous (deoxygenated) and arterial (oxygenated) blood. Blood samples were obtained by catheterizing the femoral artery and vein with PE-50 tubing, pretreated with heparin (1000 units/ml, intravenous) to prevent coagulation. Blood was drawn using a 1 ml syringe and immediately placed inside a 3D printed holder designed to fit within a 35 mm volume transceiver. Low resolution SORDINO data were acquired with: TR = 0.625 ms, FA = 3°, receiver bandwidth = 62.5 kHz, spokes = 3200, matrix size = 64^3^ and FOV = 25.6 x 25.6 x 64 mm. High resolution SORDINO data were acquired with: TR = 1.2 ms, FA = 3°, receiver bandwidth = 73.5 kHz, spokes = 80892, matrix size = 160^3^ and FOV = 25.6 x 25.6 x 64 mm. EPI was acquired with: TR = 2000 ms, TE = 14 ms, FA = 70°, receiver bandwidth = 250 kHz, matrix size = 64^2^, FOV = 40 mm^2^ and a single 1 mm thick slice. RARE images were acquired with: TR = 2500 ms, TE = 33 ms, receiver bandwidth = 54.65 kHz, matrix size = 256, FOV = 40 mm^2^ with a single 1 mm thick slice. In analysis, the signal intensities were normalized to the arterial blood signal intensity. A two-tailed unpaired t-test was used to compare the effect of blood source on signal intensity.

#### *Assessment of* receiver bandwidth and undersampling factor on SORDINO spatial resolution and noise

We further investigated how characteristics of the SORDINO sequence impact the spatial resolution and noise of reconstructed images (**Figure S8**). A defining property of SORDINO is its efficient k-space sampling by maintaining a constant total gradient angular change, enabling an extremely low acquisition bandwidth – a strategy that increases SNR by decreasing noise variance. However, as a trade-off, a longer readout may introduce spatial blurring due to signal attenuation from T2* decay during acquisition.

Another factor that may affect the spatial resolution of SORDINO images is undersampling of k-space. Like all MRI sequences, SORDINO requires a balance between sufficient temporal (∼ 2s) and spatial (∼ 0.4 - 0.6 mm^3^) resolution for rodent functional brain mapping. To achieve this balance, SORDINO data is undersampled in k-space. However, undersampling may also introduce noise and artifacts into the reconstructed image due to violation of Nyquist’s sampling criterion. To assess how these factors impact SORDINO spatial resolution and noise, we scanned a custom-made acrylic Derenzo phantom across a wide range of receiver bandwidths and undersampling factors. The phantom featured six different hole sizes (4.7, 3.9, 3.1, 2.3, 1.5, and 1.1 mm), arranged in triangular patterns around its center and filled with deionized water. The outer diameter of the phantom was 50 mm (**Figure S8**). To evaluate the impact of imaging parameters on spatial resolution, we computed the modulation transfer function (MTF). The MTF quantifies the imaging system’s ability to preserve contrast across different spatial frequencies, essentially serving as a frequency-domain characterization of imaging resolution^129^. At higher spatial frequencies, the MTF decreases towards zero, indicating poorer definition of small structures. Conversely, higher MTF values at lower spatial frequencies indicate that the sequence effectively preserves larger structures. To quantify how imaging parameters impact noise, we measured the CNR for each hole in the phantom. SORDINO images were acquired using a 72 mm quadrature transceiver with the following imaging parameters: TR = 2 ms, flip angle = 3°, receiver bandwidth = 100 kHz and spokes = 283648, matrix size = 300^3^, FOV = 60 mm^3^, and repetitions = 4. To evaluate the effect of receiver bandwidth on the MTF and CNR, SORDINO images were acquired at four receiver bandwidths: 78, 100, 150 and 200 kHz. To assess the effect of undersampling factor on the MTF and CNR, images were acquired with undersampling factors of 1, 4, 7 and 10. Python (version 3.7) and the NumPy^111^, Statsmodels^123^ and Pingouin^125^ toolboxes were used for analysis of resolution phantom data. Line profiles were drawn through the phantom in ITK-SNAP^126^. The MTF was calculated by taking the Fourier transform of line profiles through the center of the phantom holes. The maximum resolvable frequency was determined using Nyquist’s theorem calculated as: 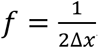, where Δ*x* is the spatial resolution. The MTF was normalized to 1 at DC, and the resolution was quantified using a threshold at 25% MTF, representing the spatial frequency at which 25% of contrast is preserved. The CNR was calculated as the difference between the mean signal within each phantom hole and the mean signal of a noise ROI, divided by the standard deviation of the noise ROI signal. The effect of imaging parameters on MTF and CNR was assessed with a linear mixed-effects model, with either receiver bandwidth or undersampling factor held as a fixed effect and each hole held as a random effect to account for the four repeated measurements.

### Rat experiments

#### Anesthetized rat MRI preparation

We followed established protocols for rat MRI procedures^23,66,70,71,130–133^. Rats were anesthetized with 4% isoflurane in atmospheric air and orally intubated with a 13-gauge catheter secured with surgical tape and sutures around the cheek. While under 2% isoflurane, PE-50 tubing lines were inserted into the lateral caudal vein using a 24-gauge catheter (Terumo, Tokyo, Japan) for anesthesia delivery. Rats were then secured in a custom-built cradle for imaging and were maintained on a sedation protocol comprising 0.5% isoflurane with a constant infusion of dexmedetomidine (0.05 mg/kg/hr) and pancuronium bromide (0.5 mg/kg/hr). Inhaled isoflurane was gradually reduced from 2% to 0.5% over the course of 30 minutes. Rats were allowed 50 minutes to establish stable physiological parameters before acquisitions began. During experiments, rats were mechanically ventilated (MRI-1 Ventilator; CWE, Ardmore, PA), end-tidal CO_2_ (ETCO_2_) was measured by a capnometer (Surgivet v9004; Smith Medical, Waukesha, WI), oxygen saturation (SpO_2_) was measured by a pulse oximetry sensor (V1703;Smith Medical, Waukesha, WI) on the foot and temperature was monitored by rectal probe and maintained by circulating warm water bath (Thermo Scientific, Waltham, MA). During experiments, ventilator flow rate, tidal volume, inspired oxygen concentration and water bath temperature were adjusted to maintain rats in a healthy and stable physiological condition (ETCO_2_ = 2.7-3.3%, SpO_2_ > 92%, and rectal temperature = 37°C).

#### SORDINO contrast in live and postmortem rat brain

To assess the SORDINO contrast in vivo, inspired by seminal experiments investigating the BOLD contrast^134^, we acquired SORDINO images of the rat brain before and immediately after sacrificing the subject with pentobarbital (120 mg/kg i.p.), followed by shutting off the ventilator inside the magnet. SORDINO data were acquired at 0.25 mm^3^ isotropic spatial resolution with the following parameters: TR = 0.62 ms, FA = 3°, spokes = 65140, receiver bandwidth = 131.58 kHz, matrix size = 144^3^, and FOV = 36 mm^3^.

#### Forepaw electrical stimulation in rats

Unilateral forepaw stimulation was delivered to the right forepaw of anesthetized rats^135–139^. Two 27 G needle electrodes were inserted between digits II and III and III and IV, then secured with surgical tape. Stimulation current accuracy was verified using a multimeter before electrode insertion. The current was delivered via a stimulus isolator (A385RC, World Precision Instruments, Sarasota, FL), triggered in synchronization with the waveform generator by a TTL signal at the start of the fMRI acquisition. The simulation parameters were set as follow for all experiments: current = 2 mA, pulse width = 0.5 ms, and frequency = 9 Hz. A block-design paradigm was employed comprising 10 s of dummy scans to allow magnetization to reach a steady-state, followed by an initial 60 s rest period and five repetitions of a 30 s stimulation period alternating with 90 s rest periods.

#### Validating TR and FA selection in SORDINO

Our Bloch equation modeling identified a range of parameters with the strongest sensitivity for fMRI. We carried out electrical stimulation of the rat forepaw using SORDINO parameter choices to validate our model at a 2 s temporal resolution and an isotropic spatial resolution of 0.6 mm^3^ in 6 rats. All images were reconstructed to an isotropic matrix size of 70^3^ and FOV of 42 mm^3^ and acquired using a homemade 1 cm ID surface transceiver. The parameter pairs examined were: (1) TR = 0.60 ms, FA = 3°, bandwidth per pixel (bw/pixel) = 905.8 Hz, spokes = 3330; (2) TR = 1.82 ms, FA = 4°, bw/pixel = 284 Hz, spokes = 1098; (3) TR = 2.44 ms, FA = 5°, bw/pixel = 208 Hz, spokes = 888; (4) TR = 2.44 ms, FA = 2°, bw/pixel = 208 Hz, spokes = 888; (5) TR = 0.60 ms, FA = 5°, bw/pixel = 905.8 Hz, spokes = 3330. Rat physiological conditions were recorded before each of the SORDINO and EPI acquisitions. Physiological parameters are expressed as mean ± SEM: SpO_2_ = 93±1 %, ETCO_2_ = 3.00±0.1 % = 37.1 ± 0.45 °C

#### Comparison of SORDINO- and EPI-fMRI sensitivity

We compared the sensitivity of SORDINO with EPI in fMRI experiments using a rat forepaw electrical stimulation model at identical temporal (2 s) and spatial (0.6 mm^3^) resolutions in 16 rats using a homemade 1cm ID surface transceiver for acquisition. SORDINO was acquired with: TR = 1.82 ms, FA = 4°, bw/pixel = 284 Hz, spokes = 1098, reconstructed matrix size = 70^3^, reconstructed FOV = 42 mm^3^. EPI was acquired with a 2D encoding trajectory: TR = 2s, TE = 14 ms, FA = 70°, bw/pixel = 5208.33 Hz, matrix size = 48^2^, FOV = 28.8 mm^3^, slices = 32 and slice thickness = 0.6 mm. Rat’s physiological conditions were recorded before each of the SORDINO and EPI acquisitions. Physiological parameters are expressed as mean ± SEM: SpO_2_ = 95±1 %, ETCO_2_ = 2.9±0.1 % = 37.7 ± 0.40 °C.

#### Data preprocessing

All SORDINO and EPI datasets underwent minimal preprocessing using the Analysis of Functional NeuroImages software suite (AFNI)^140,141^ to ensure that the evoked responses were compared with minimal confounding influences. The datasets were motion corrected using the 3dvolreg command. EPI data underwent slice-timing correction using the 3dTshift command. Next, mean images were calculated from each acquisition and skull stripped. One mean image from each subject was used to generate a study-specific template using the AntsMultivariateTempalteConstruction2 function from the Advanced Normalization Tools (ANTs) package^142^. Mean images were then aligned to the template using a series of transformations: first with rigid-body alignment, followed by affine transformation, and finally with non-linear SyN transformations using the ANTsRegistration function.

#### Analysis of SORDINO and EPI forepaw electrical stimulation data

Python (version 3.7) and the NumPy toolboxes were used for analyzing minimally preprocessed and skull stripped SORDINO and EPI data. Individual subject-level fMRI activity was analyzed using a generalized linear model (GLM) which was solved algebraically. To minimize biases from canonical hemodynamic response functions (HRFs), the HRF was empirically estimated as the average across all subjects for both SORDINO and EPI. GLM analysis incorporated regressors for a constant, first- and second-order polynomials, and a stimulation paradigm represented by four box-car function with variable delays extending up to three volumes post-stimulation onset. The beta value corresponding to the maximum response for each voxel was selected for further analysis. Active voxels were defined using a threshold of q<0.001 following false discovery rate (FDR) correction. Time courses from active voxels were subsequently extracted from the S1 forelimb (S1FL) region after removing first- and second-order polynomial trends. The same analysis was repeated for SORDINO and EPI data, replacing the box-car function with the estimated HRFs. Sensitivity was assessed by comparing z-scores, calculated as the difference between the time course signal and the mean signal intensity of the initial rest period, divided by the standard deviation of the same rest period. The CNR was defined as the average peak z-score during the stimulation period at the individual voxel level. Percent signal change was calculated similarly by subtracting the mean signal intensity of the initial-rest period from the time course, dividing by the mean rest-period intensity, and multiplying by 100. In the TR and FA selection validation experiments, the effect of parameter combinations on CNR was assessed using a linear mixed-effects model, with parameter combination as a fixed effect and subject as a random effect to account for five repeated measurements per subject. Similarly, in the sensitivity comparison experiments, a linear mixed effects model was used to assess the effect of sequence on CNR, with sequence as a fixed effect and subject as a random effect. In the head-only versus whole-body excitation experiments, the 5 repeated CNR measurements of each subject were averaged. To test whether the paired differences in percent signal change between the two coil configurations followed a normal distribution, a Shapiro-Wilk test was conducted. A paired t-test was then conducted to compare the sensitivity between the two coil configurations.

#### Comparison of SORDINO and EPI resting-state functional connectivity

A subset of nine rats from the forepaw stimulation experiments underwent 15-minute resting state scans with identical imaging parameters. All analyses were conducted on minimally processed SORDINO and EPI data. Regional Homogeneity (ReHo) analysis^143^ was performed, with mean ReHo values computed for the sensorimotor cortex and amygdala. To compare the temporal variability of FC and the spatial similarity of FC maps between SORDINO and EPI, the bilateral M1 was selected as the seed region for evaluating tSNR and FC. For tSNR calculation, the mean time series of all voxels within the bilateral M1 seed region was extracted. tSNR was computed as the ratio of the mean signal intensity to the standard deviation across time points within each temporal window. Sliding windows of varying lengths (ranging from 10 to 440 timepoints in increments of 10) were applied. For each window length, 100 randomly selected time segments were analyzed, and tSNR was computed for each windowed segment. For FC analysis, the mean time series of all voxels within the M1 seed was extracted, and the Pearson correlation coefficient was computed between the seed time series and the whole brain on a voxel-by-voxel basis. Sliding windows of varying lengths (ranging from 10 to 440 timepoints in increments of 10) were applied. For each window length, 100 randomly selected time segments were analyzed with FC values z-transformed, and their variability was estimated as the standard deviation across windows. To evaluate spatial similarity of FC, the right M1 was used as the seed region. The mean time series of the right M1 was taken as the seed, and voxel-wise Pearson correlation was performed between the seed and all other voxels within the brain. As in the analysis of the temporal variability of FC, temporal windows of varying lengths were applied and for each window length, 100 randomly selected time segments were analyzed. Spatial similarity was assessed by computing the Pearson correlation between each windowed FC map and the full-time FC map. A two-tailed paired t-test was used to assess the effect of sequence on ReHo in the amygdala.

#### SORDINO contrast mechanism in vivo: comparison of head-only and whole-body excitation

After confirming that SORDINO detects functional activations in rats with similar sensitivity to EPI, we designed experiments to test our hypothesis: that the SORDINO contrast arises from a combination of inflow-weighted CBV and tO_2_. As a first step in validating the SORDINO contrast mechanism in-vivo, we aimed to determine the fractional contributions of CBV and tO_2_ by comparing the evoked response magnitudes under two conditions: head-only and whole-body excitation. Whole-body excitation significantly reduces inflow-related vascular signal enhancement, causing all blood to reach a similar steady-state as stationary tissue. In this condition, an increase in CBV is expected to cause a modest decrease in the SORDINO signal. On the other hand, with head-only excitation, blood is excited only while flowing within the sensitivity region of the transmitter. Based on published arterial transit times^116^, blood will remain in a dynamic state, leading to an enhanced vascular signal compared to whole-body excitation. Here, an increase in CBV is expected to enhance the SORDINO signal. For both the head-only and whole-body excitation conditions, the same homemade 1cm ID surface coil functioned as a transceiver and receiver, respectively. For whole-body excitation, a 72 mm volume transmitter (Bruker Corp., Billerica, MA, USA) was activated while the surface coil was decoupled. The coil configuration was switched without repositioning the animal or the coils. By comparing the evoked response amplitude between the two conditions, we estimated the fractional contribution of CBV and tO_2_ to the SORDINO contrast. Acquisitions were performed in a randomized order in six rats using the rat SORDINO imaging parameters described above.

#### SORDINO contrast mechanism in vivo: FSCV

For in vivo FSCV experiments, polyimide/fused silica carbon-fiber microelectrodes were fabricated following established methods^48,62,144^. Briefly, a 7 µm-diameter carbon fiber (Thornel T-650) was threaded through a silica/polyimide capillary (#1,068,150,381; Polymicro Technologies Inc., Phoenix, AZ, USA) and sealed with clear epoxy. Approximately 100 µm of carbon fiber protruded from one end of the capillary to serve as the active surface area. On the opposite end, carbon fibers were adhered to a silver connection wire using consecutive layers of silver epoxy, silver paint, and clear epoxy. The electrodes were chronically implanted to the ventral striatum (coordinates: 1.3 mm A-P, 2.0 mm M-L, 6.8–7.8 mm D-V relative to bregma) 4 days prior to the experimental data acquisition to ensure post-surgical recovery. In addition to the carbon fiber electrodes, an MRI-compatible guide cannula (BASi Research Products) was implanted in the opposite hemisphere. The guide cannula was used for placing a freshly chloridized Ag/AgCl reference electrode on the day of FSCV recording. For FSCV experiments in MRI, data were acquired using High-Definition Cyclic Voltammetry and Analysis software, respectively (UNC Electronics Facility, University of North Carolina at Chapel Hill, Chapel Hill, NC). Custom instrumentation from the UNC Electronics Facility was used, including an MRI-compatible headstage connected to the chronically implanted electrodes via a triaxial cable. An oxygen-sensitive voltage waveform^145^ was scanned at the electrode surface at 400 V/s and a rate of 10 Hz^63^. The waveform first scans from 0 V to +0.8 V to oxidize DA, then down to –1.4 V to reduce molecular tissue oxygen, and finally returns to 0 V (**Figure 3e**). Current versus time traces recorded at the reduction currents of the analytes of interest were extracted and converted to concentration using fiber length-normalized *in vitro* calibration factors acquired previously (–0.19 nA/uM for oxygen^48^) at fused silica capillary fiber microelectrodes.

#### SORDINO contrast mechanism in vivo: spectral fiber photometry

An established spectral fiber photometry system was used^60,71,72,146,147^. A 561 nm CW laser (OBIS 561 LS-50, Coherent, Santa Clara, CA) was directed into a fluorescence cube (DFM1, Thorlabs, Newton, NJ). Neutral density filters (NEK01, Thorlabs, Newton, NJ) placed before the fluorescence cube adjusted laser power. The fluorescence cube incorporated a dichroic mirror (ZT488/561rpc, Chroma Technology Corp) to reflect and direct the laser beam through an achromatic fiber port (PAFA-X-4-A, Thorlabs, Newton, NJ) into a 105/125 μm core/cladding multi-mode optical fiber patch cable. The distal end of this cable connected to an implanted optical fiber probe, enabling both excitation laser delivery and emission fluorescence collection. Emission fluorescence traveled back along the patch cable to the fluorescence cube, passed through the dichroic mirror and an emission filter (ZET488/561m, Chroma Technology Corp, Bellows Falls, VT), and was directed via an aspheric fiber port (PAF-SMA-11-A, Thorlabs, Newton, NJ) into an AR-coated 200/230 μm core/cladding multi-mode patch cable (M200L02S-A, Thorlabs, Newton, NJ). This cable delivered the fluorescence to a spectrometer (QE Pro-FL, Ocean Optics, Largo, FL) for spectral data acquisition, controlled by OceanView software (Ocean Optics, Largo, FL). To minimize background light, the scan room was darkened before each session, and laser power was adjusted to balance spectral amplitudes (maximum <100 μW). A background spectrum was recorded and subtracted during acquisition. Spectral fiber-photometry data were collected at 10 Hz. Using an established workflow in MATLAB (R2024b, MathWorks)^60,146^, the TdTomato emission spectrum was processed to derive HbT, HbO, and HbR based on known molar extinction coefficients, photon pathlengths from Monte Carlo simulations, and spectral data. The generalized method of moments (GMM) was used to calculate molar concentration changes for HbO and HbR, with HbT determined as their sum. HbO and HbT changes were Z-normalized for group-level analyses.

#### SORDINO contrast mechanism in vivo: concurrent SORDINO-FSCV and SORDINO-photometry

We aimed to measure SORDINO signals during gas-challenge experiments alongside ground-truth photometry measurement of hemodynamic parameters and CBV, as well as FSCV measurement of tO_2_. The experiments were designed dissociate changes in tO_2_ from CBV. Four rats underwent simultaneous SORDINO acquisitions and spectral fiber photometry, while three rats underwent simultaneous SORDINO and FSCV recordings. All gas-challenge trials followed a block design, consisting of 300 s of baseline gas inhalation, followed by 300 s of a gas challenge, repeated three times. SORDINO imaging parameters were: TR = 1.79, flip angle = 4°, spokes = 1098, bw/pixel = 309 Hz, reconstructed matrix size = 70^3^, and reconstructed FOV = 42 mm^3^. Rats were prepared for MRI as outlined in the Rat fMRI section, with differences described in the relevant sections.

In Spectral Fiber Photometry, changes in total blood hemoglobin concentration (HbT, a.k.a. CBV), oxyhemoglobin (HbO), and deoxyhemoglobin (HbR) were computed using our previously published spectral fiber-photometry approach^60^. TdTomato was expressed under the CAG promoter in the rat S1 (coordinates: AP: 1 mm, ML: 4 mm, DV: 0.9 mm) via AAV (59462-AAV9, Addgene, Watertown, MA) microinjection, followed by a 3-week incubation for viral expression. Concurrent SORDINO-fMRI and spectral fiber-photometry recordings during a gas-challenge were performed (**Figure 3d**). The experiment investigated the hemodynamic response to different inhaled gas conditions. Initially, rats were ventilated with a hypercapnic-hyperoxic mixture (10% CO_2_ and 90% O_2_), which was expected to cause vasodilation compared to when breathing atmospheric air. During the gas challenge, the inhaled mixture was switched to a hypercapnic-hypoxic mixture (10% CO_2_ and 10% O_2_), which we expected to reduce tO_2_ and further increase CBV due to hypoxia-induced vasodilation^148^.

In FSCV, a silica-based carbon-fiber electrode was chronically implanted into the ventral striatum. The voltammetric electrodes were connected to an MRI-compatible FSCV headstage via a triaxial cable^48^, and the animal was positioned inside the MRI scanner. An N-shaped waveform was applied to detect changes in tissue oxygen levels. This oxygen-sensing waveform consisted of an 11 ms scan holding at 0 V, ramping up to +0.8 V, descending to −1.4 V, and returning to 0 V at a scan rate of 400 V/s versus Ag/AgCl^62,63,145^. The waveform was applied at 10 Hz, with data sampled at 100 kHz. Two different gas-challenge experiments were carried out with simultaneous SORDINO and FSCV. Each experiment was designed to elicit an opposite change in tO_2_ – one an increase and another a decrease. In the first experiment, rats initially inhaled compressed room air (21% O_2_) as the baseline, before transitioning to a hyperoxic mixture (100% O_2_) to induce an increase in tO_2_. In the second experiment, rats first inhaled a hypercapnic-hyperoxic baseline (10% CO_2_ and 90% O_2_) before switching to a hypercapnic-hypoxic mixture (10% CO_2_ and 10% O_2_) to induce a decrease in tO_2_.

SORDINO data were reconstructed using an in-house tool incorporating the *mri-nufft*^149^ Python module and *finufft*^150,151^ as the backend. All data were analyzed in MATLAB (R2024a). Time courses from a voxel at the tip of the fiber-photometry fibers were converted to z-scores by subtracting the mean signal intensity of the baseline period from the time course and dividing by the standard deviation of the same baseline period. Simultaneously collected FSCV and SORDINO data were normalized to the mean of 9 trials from 3 subjects. All time courses were aligned to a time corresponding to the beginning of the gas challenge paradigm.

### Mouse experiments

#### Awake mouse fMRI preparation – implantable headplate coil

We developed a head-fixed setup for awake mouse imaging using a sterile, implantable PCB headplate designed for both head stabilization and as a transceiver for imaging. The headplate, measuring 1.2 x 2.8 x 1.5 cm, featured a 0.9 cm circular opening at the center for recording probe implantation. The PCB incorporated copper circuit traces (short axis 15, long axis 15 cm and a 1mm trace width with a thickness of 70 µM) pre-printed on both sides and insulated by a 10 µM soldermask layer. This headplate coil functioned as an RF transceiver or receiver, depending on the resonance circuit board connected. As a head-fixation system, the headplate locked into two securing plates attached to the animal cradle and interfaced with a custom resonance circuit board via a 2-pin socket connection.

#### Awake mouse fMRI preparation – headplate implantation procedures

The implantation was performed during the same surgical session as the virus injections and GRIN lens implantation. Mice were anesthetized with isoflurane (1-1.5 %) in medical air. The head was stabilized in a stereotaxic frame, and if no existing incision from a prior procedure was present local anesthesia (0.1 mL, 0.5% bupivacaine subcutaneous) or lidocaine spray was applied before carefully removing a piece of scalp (< 1 x 1 cm) to expose the skull. The headplate was secured trace-side down using a C&B Metabond Quick Adhesive Cement (Parkell, Brentwood, NY). The skull was cleaned with Bactine followed by application of dental etching gel for 5 – 10 minutes and then rinsed again with Bactine. The headplate was then positioned and adhesive cement was applied to seal the incision and secure the headplate to the skull. Post-surgical care comprised saline (1.5 ml, subcutaneous) for hydration and access to hydrogel inside the home cage. Meloxicam (0.1 mg/kg, oral) was provided for 2-3 days post-surgery as needed. Lincomycin hydrochloride (30 mg/kg, intramuscular) was administered to prevent infection. Mice were allowed at least one week to recover before further experimental procedures.

#### Awake mouse fMRI setup and SORDINO parameters

The headplate coil was mounted directly atop the mouse skull as described above. The mouse was positioned on a custom-built cradle, and the headplate coil was connected to a circuit board. Once secured to the holder, the mouse was allowed to walk freely on a custom-built, MR-compatible treadmill, a strategy employed in head-fixed two-photon imaging studies to reduce stress-related confounds^152,153^. A high resolution SORDINO image was acquired as a reference with the following parameters: TR = 5 ms, FA = 7°, spokes = 80876, receiver bandwidth = 75 kHz, matrix size = 160^3^, and FOV = 25.6^3^. Functional imaging with SORDINO was performed using the following parameters: TR = 0.62 ms, FA = 3°, spokes = 3200, receiver bandwidth = 60 kHz, matrix size = 64^3^, and FOV = 25.6^3^. This acquisition is referred to as the default SORDINO mouse fMRI parameters throughout the study.

#### Serum cortisol level

To evaluate stress in awake mice undergoing SORDINO and EPI, serum cortisol levels were measured during habituation (**Figure 2b**). Mice were divided into 3 cohorts: 32 mice undergoing habituation with SORDINO, 6 mice with EPI, and an age-matched baseline control group (49 mice) corresponding to day 1 of habituation. Serum samples were collected from the submandibular gland of mice on days 1, 3, and 5 using Microtainer Tubes with Serum Separator Additive and Microgard Closure (BD, MFR# 365967). Blood was allowed to clot at room temperature for 30 minutes, followed by centrifugation at 2,000 × g for 10 minutes to separate the serum, which was then stored at −80°C until analysis. Cortisol levels were quantified using the Cortisol Parameter Assay Kit (KGE008B, R&D Systems, Bio-Techne, Minneapolis, MN, USA) following the manufacturer’s protocol. Absorbance was measured at 450 nm, and cortisol concentrations were calculated using a standard curve. A one-way ANOVA tested the effect of sequence on serum cortisol levels, followed by Tukey’s HSD for post-hoc comparisons.

#### Body Weight

The weight of a subset of 15 and 6 C57BL/6J mice undergoing habituation with SORDINO and EPI respectively were measured on days 1, 3 and 5 of habituation (**Figure 2c)**. A repeated measures ANOVA was used to test the effect of habituation day on body weight. Tukey’s HSD was used for post-hoc comparisons.

#### Miniscope calcium imaging

We performed calcium imaging in an awake mouse with an in-house modified MRI-compatible miniscope system (base model: nVista system, Inscopix, Mountain View, CA) (**Figure 2g-h**). A ProView™ Integrated GRIN lens (Inscopix, Mountain View, CA) with 0.6 mm diameter was surgically implanted into the medial prefrontal cortex (AP = 2.0 mm, ML = 0.35 mm, DV = −2.1 mm), positioned approximately 0.1 mm above the virus injection site, following AAV-CaMKIIα-jGCaMP8m injection. The virus was injected at a volume of 1 μL at a flow rate of 0.1 μL/min, with an additional 10-minute diffusion period before slow retraction of the microsyringe needle. Animal preparation was as described in awake mouse fMRI and imaging was acquired using the default mouse SORDINO imaging parameters. Data were recorded over a 10-minute period, with equal durations of scanner idle state and SORDINO acquisition. Raw calcium miniscope data were preprocessed using Inscopix Data Processing Software (IDPS), which included motion correction, spatial filtering, ΔF/F normalization, and PCA/ICA for neuronal component identification.

#### Assessment of motion artifacts

The sensitivity of SORDINO and EPI to motion was evaluated in two experiments. In the first experiment, the impact of motion on SORDINO and EPI data was compared in an anesthetized mouse using controlled motion. The setup for MRI was as described in *Awake mouse fMRI* above, except that the mouse was anesthetized with pentobarbital (50 mg/kg, intraperitoneal). A nylon monofilament string was tied around the right forepaw, allowing controlled movement from the console workstation during scanning. The experiment followed a block design, alternating between 10 s of rest and 10 s of controlled forepaw movement at a frequency of 1 Hz, repeated nine times. In the second experiment, thirty awake mice underwent both SORDINO and EPI scans for motion assessment. Images from each timeseries were aligned to the first volume, which was used as the reference using AFNI’s 3dvolreg command. This command generated six motion parameters: three for rotation (roll, pitch, and yaw) and three for displacement (Inferior-Superior, Left-Right, and Anterior-Posterior). Python (version 3.7) with NumPy^111^, MATLAB (R2024a) and GraphPad Prism 10 were used for data analysis.

To compare the effect of SORDINO and EPI on motion parameters, motion for each subject was quantified as the mean absolute value of rotation and displacement across the time series. Paired t-tests were performed to compare the effect of imaging sequence on each of the six motion parameters. In the controlled motion experiment, Pearson correlation coefficient was calculated between the brain signal time courses and the movement blocks for both SORDINO and EPI data on a voxel-by-voxel basis. The correlation coefficient values underwent Fisher’s Z-transformation and results were visualized on a histogram.

#### Awake mouse resting-state fMRI experiment

A total of 25 well-habituated male C57BL/6J mice, weighing 28-32 g, were used in this study. Functional (30 minutes per session) and high resolution anatomical SORDINO images were acquired using the default mouse SORDINO imaging parameters and reconstructed using an in-house tool that incorporates the *mri-nufft*^149^ Python module with *finufft*^150,151^ as the backend. Brain masks for skull stripping were manually drawn for each subject using ITK-SNAP. Spatial normalization was performed to generate a population average brain template. To ensure alignment to a common framework, the Allen Mouse Brain Common Coordinate Framework (CCF) image, including its annotated regions of interest (ROIs), was realigned to RAS+ orientation and centered at the anterior commissure. Subsequently, the population average brain template was registered to the AC-centered Allen Mouse Brain Template. Functional SORDINO reconstruction involved spoke timing correction applied to the FID signal, followed by a NUFFT. Motion correction was performed using frame-wise displacement, calculated based on a 5 mm radius for the mouse brain. The functional images were co-registered to the population average brain template. Signal processing steps included nuisance regression of 28 parameters: a third-order polynomial (4), six rigid motion parameters (6), one TR time shift and its squared terms (12), along with their squared terms. Temporal filtering was applied using a band-pass filter with a frequency range of 0.01–0.15 Hz. Spatial smoothing was performed using a Gaussian kernel with a full width at half maximum (FWHM) of 0.6 mm, corresponding to 1.5 times the voxel size (0.4 mm³).

#### Assessment of awake mouse resting-state fMRI quality

To ensure data quality, we performed multiple quality control assessments, including FD, DVARS, tSNR, and FC specificity measures^35,66,154^ (**Figure 4b-e**). *FD and DVARS*: FD was calculated to evaluate head motion, with a threshold of 0.08 mm used to identify and censor frames with excessive movement. DVARS, a well-established metric, was calculated to measure the rate of change in the BOLD signal intensity between consecutive time points, serving as an additional check on signal stability. *tSNR:* The tSNR was computed as the ratio of mean signal intensity to the standard deviation of the noise over time for each voxel. This metric evaluates the signal quality across the dataset, where higher tSNR values indicative of better data quality. tSNR was initially high but decreased before stabilizing, likely due to increasing physiological noise, minor head motion, and slow BOLD signal drift over time^155^. *FC Specificity*: As part of the quality control process, FC specificity was assessed using a non-spatial Gaussian/Gamma mixture modeling approach^156^ to determine the threshold for connectivity metrics. Voxel-wise connectivity maps derived from seed-based analyses of the S1L and S1R, were used to create a sample distribution. The data were Fisher Z-transformed and standardized before the mixture modeling. A probability map was generated across a range of z-values, where voxels with low connectivity to the seed region were modeled by the Gaussian component (blue), and voxels with high connectivity to the seed region were captured by the Gamma component (red). The intersection point between these distributions was identified and used as the threshold for defining functionally specific connectivity.

#### Mapping intrinsic mouse brain functional connectivity networks

To identify intrinsic brain functional connectivity networks from awake SORDINO-fMRI data, group-level ICA was performed using the MELODIC tool from the FSL software package^157^. ICA analysis was conducted with the number of components fixed at 20. To threshold the ICA spatiotemporal maps, a Gaussian/Gamma mixture model was applied to the histogram of intensity values using the Expectation-Maximization (EM) algorithm implemented in MELODIC. The resulting z-score maps of the ICA components were thresholded at a probability level >0.3, facilitating the identification of significant independent components representing intrinsic functional connectivity networks. Among the identified ICA components, the connectivity structure representing the Triple Networks, including the DMN, LCN, and SN, was selected as a representative map due to its comparable spatial patterns. This selection highlights the intrinsic brain functional connectivity networks identified in the dataset.

#### Pair-wise ROI functional connectivity matrix and clustering

The Allen Institute’s SDK was used to provide the mouse brain reference template space and anatomical connectivity data via their Mouse Connectivity Database. ROIs and atlas images were generated using structural set ID 687527945 with the AllenSDK. The atlas image was aligned to the population average template and downsampled to a voxel size of 400 μm³. To ensure robustness, only ROIs larger than 4 voxels were retained, resulting in a total of 148 ROIs for analysis. Functional connectivity between the 148 ROIs was estimated using pairwise Pearson correlation coefficients. The resulting correlation matrix was Fiser Z-transformed to improve normality. A permutation-based one-sample t-test with 5000 permutations was conducted to statistically evaluate the functional connectivity matrix. Significance was assessed at p < 0.05 (two-sided), using the distribution of minimum and maximum t-values derived from the permuted datasets. To identify functional modules, the group-level functional connectivity matrix derived from the permutation t-test was thresholded at <0.1 to remove low connectivity edges. The thresholded matrix was then converted into a weighted graph. The Louvain community detection algorithm was applied 1000 times to this graph, and the partition that maximized modularity was selected as the final modular structure. To further refine and organize the detected modules, hierarchical clustering was performed, using Ward’s hierarchical clustering method. A dendrogram was constructed, and thresholds were iteratively applied. The number of modules identified by the Louvain community detection algorithm was used as the minimal partition, while higher thresholds were applied to generate hierarchical organization and assign cluster labels. The same analysis pipeline used for functional connectivity was applied to the anatomical connectivity matrix to identify anatomical modules, ensuring consistency between clustering approaches for functional and anatomical data. To integrate these results, a combined classifier was generated by concatenating functional and anatomical cluster labels assigned to each ROI. This multi-modal classifier provided a comprehensive representation of the connectivity features for each ROI. To ensure uniform scaling of features, the combined classifier data were standardized. Principal Component Analysis (PCA) was then employed to reduce the dimensionality of the dataset while retaining 99% of its variance, facilitating the clustering process without significant information loss. The DBSCAN clustering algorithm was utilized to classify the PCA-transformed data. To determine the optimal ɛ (epsilon) value, which defines the maximum allowable distance between points within a cluster, a k-distance graph was generated by estimating the distance to the kth nearest neighbor for each point. The KneeLocator algorithm was applied to identify the “elbow point” on the graph, which corresponds to the optimal ɛ value. Using the optimal ɛ and setting a minimum cluster size of k = 3 ROIs, DBSCAN was applied to assign cluster labels to each ROI, representing their membership in specific functional and anatomical clusters. The resulting clusters were systematically sorted and organized for subsequent interpretation and analysis.

#### Mapping the reach-and-grasp task using SORDINO-fMRI in mice

Male VGAT-ChR2-EYFP mice (C57BL/6J background, n = 4), weighing 23–25 g, were used in this study. To leverage SORDINO’s insensitivity to motion-induced artifacts and negligible acoustic noise, we trained mice to execute a forelimb reaching task simultaneously with whole-brain mapping with SORDINO. Following recovery from headplate implantation, four mice underwent a multi-stage food restriction protocol to facilitate training in a reach-to-grasp task. Initially, body weight was reduced by 10%, after which mice were gradually acclimated to head restraint. This acclimation involved placing the head-fixed mice in their home cage for 15 minutes per day over three consecutive days. After the initial phase of acclimation, mice were introduced to a darkened training environment, with a rotary motor-powered table. Food pellets, weighing 10 mg, were positioned at the edge of the table within licking distance. If mice did not spontaneously consume the pellets, their weight was gradually reduced to 80-85 % of its baseline, with a maximum restriction of 70%, stopping once voluntary pellet consumption was observed. At this stage, pellets were presented directly in front of the mouth using blunt forceps. This process was repeated until the mouse reached for and grabbed the pellet from the forceps. Once the association was established, pellets were again placed on the rotary table but positioned beyond licking distance to encourage reaching behavior. Training progressed as mice gradually learned to grab pellets from the table. Mice were then trained to associate an auditory cue of 5 kHz frequency and 500 ms duration with food delivery. Pellets were withheld and then presented by rotating the training table in synchrony with the auditory cue. Training continued until mice consistently reached for the pellet in response to the tone and achieved reaction times of less than approximately 250 ms. At the next stage of training, mice were transferred to a custom-built cradle designed to replicate the reach-to-grasp behavior within the MRI environment. The custom-built cradle comprised (1) a conveyor belt with slots to hold the pellets; (2) a pellet dispenser for controlled food delivery; (3) an MR-compatible speaker positioned behind the mouse for auditory cue presentation; (4) a light source made from a flexible plastic tube to transmit light from a laser; (5) a vacuum and air-line system integrated with the pellet dispenser for automated dispensing. All components of the system were controlled via an Arduino program. The cradle was attached to a motor via a pulley system, which was secured to a wooden base. During each trial, the conveyor belt rotated dropping a pellet into the dispenser. An auditory cue was delivered, and a small air line propelled the food pellet. Once reliable performance was achieved, a back cover was added to the cradle to prevent mice from arching their back and neck towards the headplate. Training continued until mice reached sufficient accuracy, typically achieving 65 – 90% success rates. In the final phase of training, mice were gradually acclimated to the MRI scanner environment. Initially, the cradle was placed outside the scanner. Mice were trained, as previously described, in this location for 3 consecutive days.

Then, mice were placed inside the magnet bore and training continued. Scanning sessions commenced once mice achieved a success rate more than 50%. This final stage of training typically required an additional 3-7 days. Each fMRI session, using the default mouse SORDINO imaging parameters, comprised 75-150 reaching trials, with each trial lasting 20 s. At the start of each trial the pellet was dispensed with the auditory cue. The reaching movements were recorded at 30 Hz using an MR-compatible camera. To maintain precise alignment throughout the recording, the light source was briefly flashed off for 100 ms before being turned back on. The flash served as a timestamp to correct for potential drift of the Arduino clock. The Arduino microcontroller was programmed to send a TTL pulse to the MRI scanner every 2 s, triggering the acquisition of a 1.987 s volume, ensuring that each volume was precisely synchronized with the Arduino trigger. The pellet dispensing and auditory cue presentation occurred at the start of the corresponding volume, ensuring that reaching movements consistently aligned with the beginning of each volume, minimizing variability in movement timing relative to volume acquisition. In analysis, the timeseries were averaged and extracted from four regions of interest: RSC, M1, S1, and Thalamus. A linear mixed-effects model was used to assess the signal changes at time points 0, 2, and 4 s after pellet dispensing with the mean baseline signal change averaged over 2, 4, and 6 s before pellet dispensing. To account for repeated measurements of the same subject, subject included as a random effect. A threshold was set at an FDR-corrected *p* < 0.05.

#### Mapping inter-brain synchronization in socially interacting mice

Male C57BL/6J mice (n = 22), weighing 25–28 g, were used in this study. The mice were grouped into 11 pairs and underwent hyperscanning with SORDINO. Mice were prepared as described above, except that a custom-designed 3D-printed cradle was used to position two awake mice face-to-face, separated by a remotely retractable divider. Each mouse was equipped with a head-mounted surface RF transceiver coil, enabling simultaneous dual-brain imaging during social interaction. Mice were habituated to the setup over five consecutive days before scanning. During hyperscanning, mice were head-fixed face-to-face with a preinstalled divider before being placed in the scanner. Mice underwent a 40-minute continuous recording using the default mouse SORDINO imaging parameters, comprising 20 minutes pre- and post-divider removal. For all analyses, ROIs were defined using the AllenSDK (structural set ID 687527945) following the same approach described in the *Pair-wise ROI functional connectivity matrix and clustering* section. To investigate evoked responses to social encounters, we performed peri-event analysis on the SORDINO-fMRI data. Voxel-wise SORDINO-fMRI time courses were normalized to percentage signal changes, averaged across subjects, and thresholded at 0.6% to visualize activation and deactivation time-locked to divider removal. Based on these results, we conducted peri-event analysis using SORDINO signals extracted from selected ROIs, which were normalized to the pre-divider removal baseline using Fisher’s Z-transformation. Mirrored inter-brain connectivity was assessed by computing voxel-wise Pearson correlation coefficients between spatially corresponding voxels in the two interacting brains. A one-sample *t*-test was performed to generate a parametric map, with significance assessed at *p* < 0.01. Inter-brain cross-region connectivity was evaluated by computing Pearson correlation coefficients among the ROIs from the two interacting subjects. A one-sample *t*-test was then conducted for each inter-brain ROI pair, with significance assessed at *p* < 0.001 (FDR-corrected for multiple comparisons). Spatial correlation within the DMN and SN was computed for each scan volume (500 volumes pre- and post-divider removal). Distributions of spatial were analyzed across scan volumes and subjects. To examine inter-brain connectivity to specific brain regions in the interacting partner, the SORDINO signal time course from a selected ROI in one brain was extracted and used to compute a voxel-wise Pearson correlation map in the partner’s brain. The resulting correlation maps for each subject were standardized using Fisher’s Z-transformation, followed by a paired *t*-test (post-divider removal v.s. pre-divider removal) to generate a parametric map, with significance assessed at *p* < 0.001.

#### Mapping glymphatic dynamics in mice

Male (n = 4) and female (n = 1) C57BL/6J mice, weighing 27–30 g, were used in this study. Animals were anesthetized with isoflurane (4% induction, 1–2% maintenance) and maintained at physiological body temperature using a rectal temperature probe and heat lamp throughout the procedure. Hair over the lower skull and neck was removed using Nair, and the surgical site was sterilized with alcohol pads. A midline incision was made along the neck to expose the atlanto-occipital membrane by carefully dissecting through two layers of muscle. A borosilicate glass pipette, pulled using a micropipette puller and blunted for safe use, was inserted perpendicularly through the atlanto-occipital membrane into the cisterna magna, with proper placement confirmed by a reduction in resistance. The pipette was secured in place with Vetbond and superglue to prevent movement during imaging. An intraperitoneal catheter was inserted to administer a mixture of ketamine (100 mg/kg) and xylazine (15 mg/kg) anesthesia during the scanning session. The mouse was positioned on a custom sled and carefully transferred into the 9.4T MRI scanner. Two capillary tubes containing gadoteridol (Prohence, 0.067 mM) and saline were placed alongside the animal to serve as reference phantoms for the imaging session. Dynamic contrast-enhanced SORDINO-fMRI was performed to evaluate glymphatic dynamics by imaging the influx of Gd contrast agent into the brain parenchyma. SORDINO images were acquired using the following parameters: scan time per volume = 1.5 minutes, TR = 1.1 ms, flip angle = 6°, receiver bandwidth = 78.1 kHz, spokes = 80892, matrix size = 160^3^, field of view (FOV) = 25.6^3^ mm. A baseline M0 scan was acquired for 10.5 minutes (7 volumes), followed by dynamic imaging, initiated immediately after administering 10 µL gadoteridol (Prohence, 0.067 mM) with a 2 µL saline chase via intra-cisterna magna injection using an MRI-compatible syringe pump at a rate of 0.5 µL/min. Anesthesia was transitioned from isoflurane to ketamine/xylazine via bolus injections administered through the intraperitoneal catheter, guided by physiological monitoring of respiratory rate and oxygen saturation. The total imaging session lasted 100.5 minutes. The acquired DCE-SORDINO data were reconstructed using an in-house non-uniform Fourier transform (NUFFT)-based pipeline with spoke timing correction applied to the k-space data. Motion correction was applied to reduce head motion artifacts. Reconstructed images were coregistered to a well-oriented subject dataset using SyN diffeomorphic non-linear registration implemented in the ANTs software for cross-subject alignment. Signal intensity changes were extracted from predefined ROIs, including the cerebral cortex, lateral ventricles, third ventricle, fourth ventricle, and cisterna magna, using binary masks aligned with the registered data. Dynamic contrast signals were normalized using a two-point normalization approach, leveraging the Gd and saline reference intensities obtained from the capillary tube phantoms.

#### Manganese-enhanced SORDINO

Male C57BL/6J mice (n = 3), weighing 25–26 g, were used in this study. To ensure MRI compatibility, the metallic flow moderator in osmotic mini-pumps (ALZET 1007D; DURECT, Cupertino, CA, USA) was replaced with a PEEK alternative. The pumps, with a release rate of 0.5 µL/h for 7 days, were implanted subcutaneously. Each pump contained 100 µL of 3.45 µM/µL MnCl_2_·4H_2_O (Sigma-Aldrich, St. Louis, MO) in 100 mmol bicine buffer (Sigma-Aldrich, St. Louis, MO). Following 7 days of continuous Mn infusion, reaching a total dose of 175 mg/kg, the mice underwent MRI. SORDINO images (**Figure S6b**) from the same mice, taken before and 7 days after subcutaneous osmotic mini-pump infusion of MnCl_2_ at a total dose of 175 mg/kg, were compared^89^.

## Acknowledgements

We thank Dr. Olli Gröhn at the University of Eastern Finland, Dr. Ali Özen at the University of Freiburg, and members at the UNC Center for Animal MRI for insightful discussions. This work was supported in part by the National Institute of Biomedical Imaging and Bioengineering (R01EB033790 to Y.-Y.I.S., S.-H..L., L.-M.H., and W.-T.C.), National Institute of Mental Health (RF1MH117053, R01MH126518, R01MH111429, S10MH124745 to Y.-Y.I.S.), National Institute of Neurological Disorders and Stroke (R01NS091236 and R21NS133913 to Y.-Y.I.S.), National Institute on Alcohol Abuse and Alcoholism (P60AA011605 and U01AA020023 to Y.-Y.I.S. and S.H.L.), National Institute of Drug Abuse (R21DA057503 to L.-M.H. and Y.-Y.I.S.), National Institute of Child Health and Human Development (P50HD103573 to M.D.S., B.D.P., S.S.O., Y.-Y.I.S. and S.H.L.; T32HD040127 to S.T.A.), National Institute of Health Office of the Director (S10OD026796 to Y.-Y.I.S.), and W.M. Keck Foundation (to A.W.H., S.T.A., Y.-Y.I.S., and S.S.).

## Author contributions Statement

Conceptualization, M.J.M., Y.M., W.-T.C., and Y.-Y.I.S.; Methodology, M.J.M., T.-H.H.C., S.S., L.-M.H., S.T.A., Y.M., T.A.S., S.S.O., U.E.E., M.D.S., B.D.P., A.W.H., S.-H.L., W.-T.C., and Y.-Y.I.S.; Investigation, M.J.M., S.S., T.-H.H.C., L.-M.H., S.T.A., T.A.S. T.-W.W.W., R.J.N., C.D.F., S.S.O., S.-H.L., W.-T.C., and Y.-Y.I.S.; Resources, M.J.M., S.S., T.-H.H.C., L.-M.S., S.T.A., T.A.S., M.D.S., B.D.P., A.W.H., S.-H.L., W.-T.C., and Y.-Y.I.S.; Data Curation, M.J.M., S.S., T.-H.H.C., L.-M.S., S.T.A., Y.M., T.A.S., T.-W.W.W., and S.-H.L.; Writing – Original Draft, M.J.M. and Y.-Y.I.S.; Visualization, M.J.M., S.S., T.-H.H.C., L.-M.H., S.-H.L., and Y.-Y.I.S.; Supervision, B.D.P., A.W.H., and Y.-Y.I.S.; Project Administration, M.J.M., S.S., and Y.-Y.I.S.; Funding Acquisition, M.D.S., B.D.P., A.W.H., and Y.-Y.I.S.

## Data Availability

Processed data used for discrete hypothesis testing, bar, box, and scatter plots, and the exact test statistics derived from these data are provided in the Source Data file. All other data associated with this work, including those used for fMRI response maps and time-course plots, are publicly available on UNC Dataverse: https://doi.org/10.15139/S3/AA8XAF. Source data are provided with this paper.

## Code Availability

The codes used to perform the analyses in this study are publicly available on UNC Dataverse: https://doi.org/10.15139/S3/AA8XAF. Bruker’s policy restricts the dissemination of sequence source codes to direct peer-to-peer sharing and prohibits the use of a public repository for unrestricted access. To obtain access to this sequence, please request at https://camri.org/sordino.

