## Supplementary_Figures for "SORDINO for Silent, Sensitive, Specific, and Artifact-Resisting fMRI in awake behaving mice"

**Affiliations:**

**Author list footnotes:**

[†] These authors contributed equally

[#] Current affiliation: National Institute of Neurological Disorders and Stroke, National Institute of Health, USA

**Supplementary Figures:**

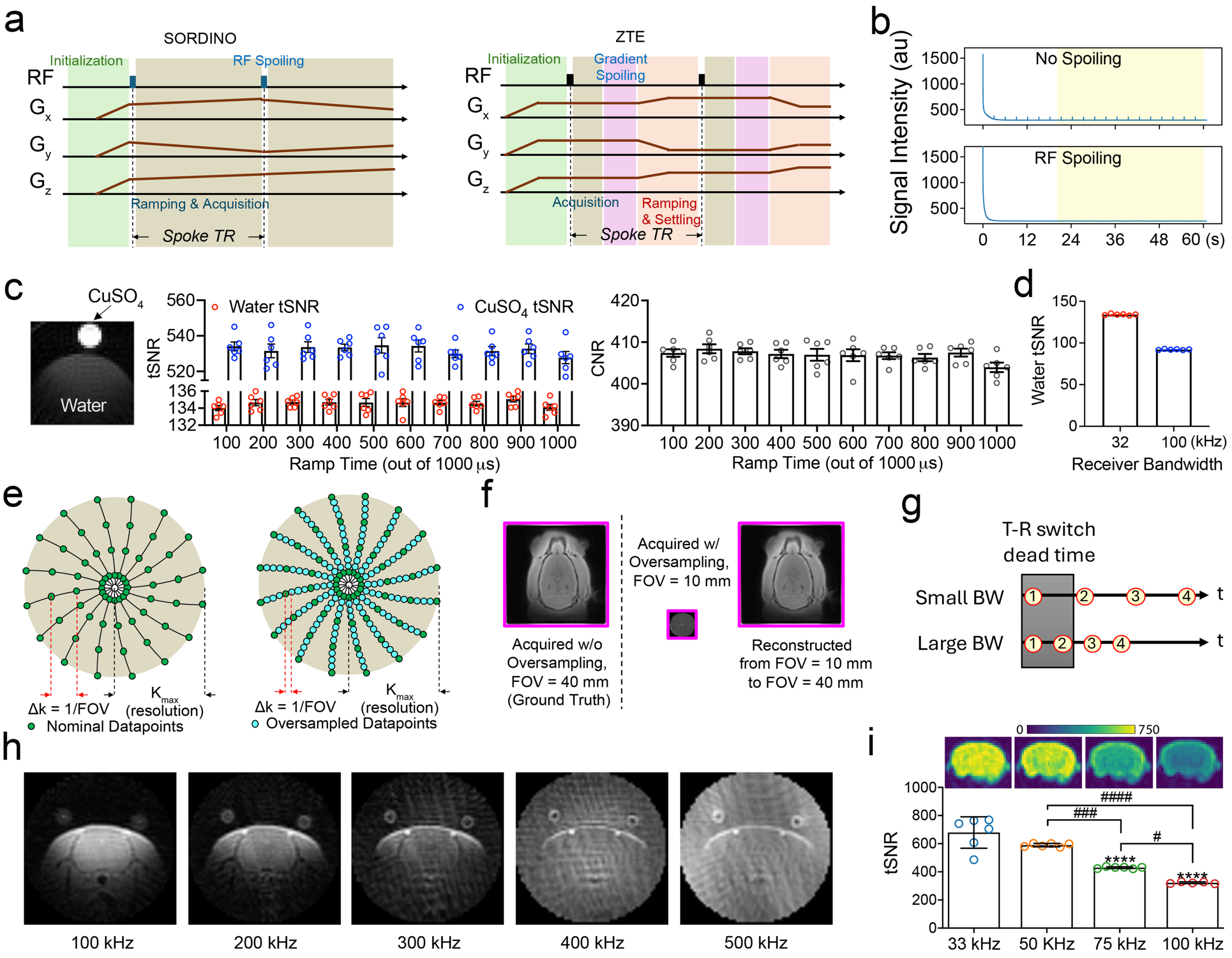


**Figure S1. SORDINO sequence and modeling of its functional contrast. (a)** Conceptual sequence diagrams showing the differences between SORDINO and ZTE. **(b)** Effect of RF-spoiling on magnetization measured from a deionized water phantom. The plots display the FID intensity evolution from a single-spoke acquisitions with and without RF-spoiling. The measured signal also validates the pattern predicted in Figure 1d. RF spoiling was implemented by incrementing the phase of the excitation pulse by 117° upon every subsequent excitation. Top: as no RF spoiling was implemented and gradient spoiling was absent, transverse magnetization was inadequately suppressed, resulting in the formation of stimulated echoes and increased temporal variability during the steady state (yellow shaded area). Bottom: RF spoiling minimized echo formation. **(c)** With identical data acquisition time, a longer ramp time does not penalize the temporal SNR (tSNR) or contrast-to-noise ratio (CNR), calculated between $CuSO_4$ and water phantoms (n = 6 repeated measurements; repeated measures ANVOA; water phantom tSNR, F(9,45) = 0.78, *p* = 0.63; $CuSO_4$ phantom tSNR, F(9,45) = 0.49, *p* = 0.87); CNR (F(9,45) = 1.28, *p* = 0.27). **(d)** With identical ramp time, a smaller bandwidth increases tSNR in the same phantom (n = 6 measurements; two-sided paired t-test, t(4) = 74.43, *p* <0.0001). **(e)** Oversampled data along the spoke direction can be used to achieve flexible FOV extension during reconstruction. The two k-space illustrations demonstrate data acquisitions reaching the same $k_{max}$ (i.e., same nominal resolution) under identical maximal gradient strengths. However, the oversampled data provided by faster analog-to-digital conversion allows FOV to be extended by any factor less than the oversampling rate. **(f)** Comparisons of rat brain images acquired with initial FOV of 40 mm versus 10 mm, with the latter also reconstructed to 40 mm using oversampled data. **(g)** A conceptual diagram illustrating the number of missing data points during a given T-R switch dead time. The numbers indicate the order of acquired k-space data points along a spoke. A smaller bandwidth reduces the number of missing data points at the k-space center, resulting in a smaller hole of empty data to be addressed during reconstruction. **(h)** At identical spatial resolution and acquisition time, brain images show improved quality at a lower acquisition bandwidth. Note that the polymeric insulation material covering the copper wire of imaging coil becomes more visible at higher acquisition bandwidths as expected. **(i)** Comparisons of lower acquisition bandwidths on brain image tSNR across multiple repeated measures (n = 6 and 5 for acquisition bandwidths 33-75 kHz and 100 kHz respectively; Ordinary one-way ANOVA followed by post hoc Tukey multiple comparison test, effect of bandwidth, $F(3, 19) = 42.26$, $p<0.0001$). Results are expressed as mean ± SEM. ****$p<0.0001$ compare with the group of 33 kHz; #$p<0.05$, ###$p<0.001$, and ####$p<0.0001$ compare with indicated groups.

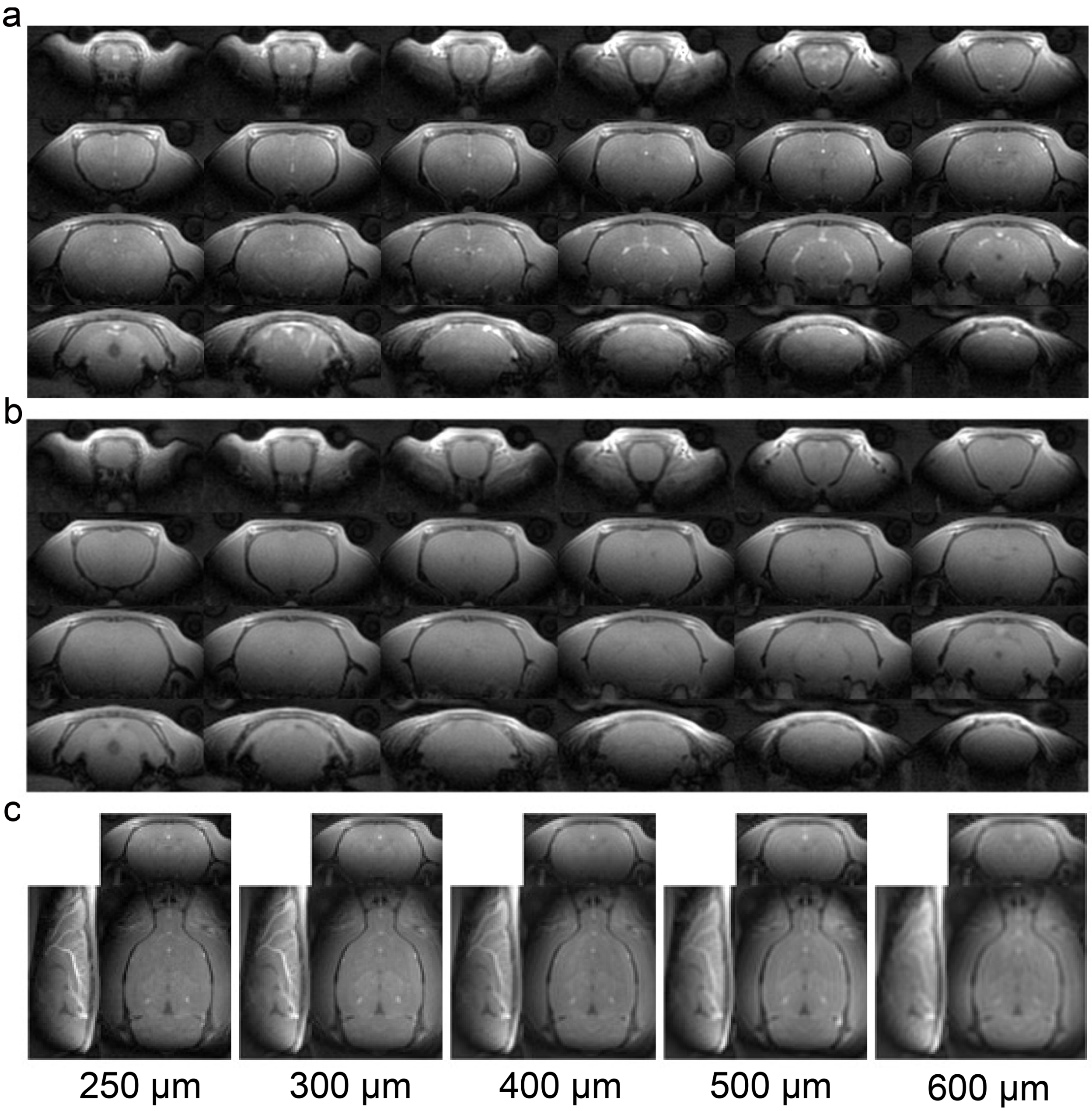


**Figure S2. Raw SORDINO images of a rat brain.** Images were acquired from a healthy adult rat using a custom-designed single-loop surface transceiver coil as seen in **(a)**, and again after euthanizing the subject in the magnet **(b)**, showing robust inflow-enhanced CBV contrast between two conditions. **(c)** In vivo SORDINO images across different isotropic spatial resolutions.

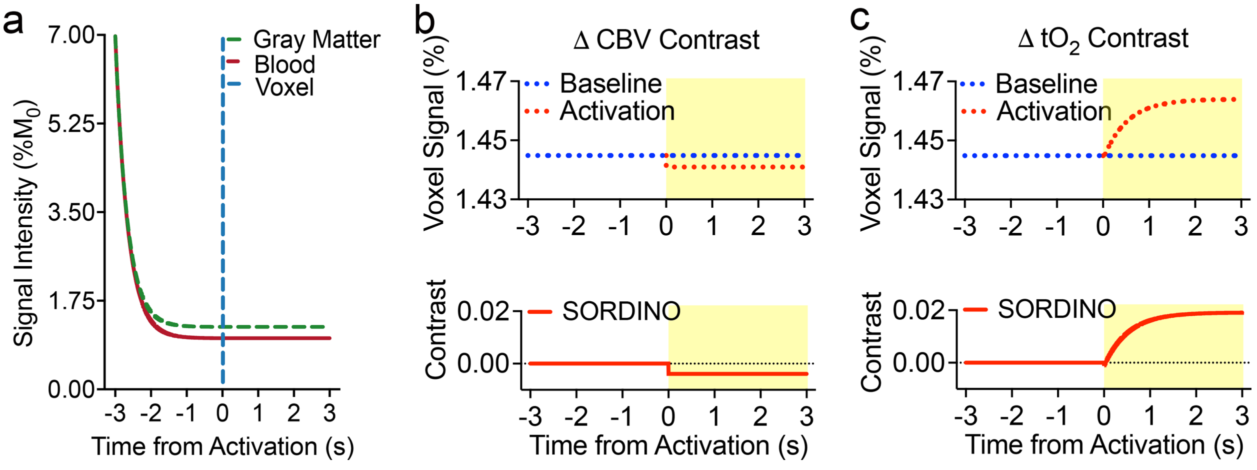


**Figure S3. Modeled SORDINO contrast using whole-body RF excitation. (a)** With a whole-body excitation coil, the measured voxel signal is comprised of a mixture of two tissue types with different T1 values. Bloch equation simulations show that magnetization of gray matter and blood reach slightly different steady-state levels within a few seconds (green dash lines). **(b)** At this steady-state, physiological increase in CBV following functional activation results in a small decrease in SORDINO signal, recapitulating the well-known VASO effect. **(c)** Physiological increase in tO2 following functional activation shortens T1 relativity and increase SORDINO signals, same as that shown in **Figure 1f**. This effect is at least an order of magnitude stronger than the weak negative CBV changes.

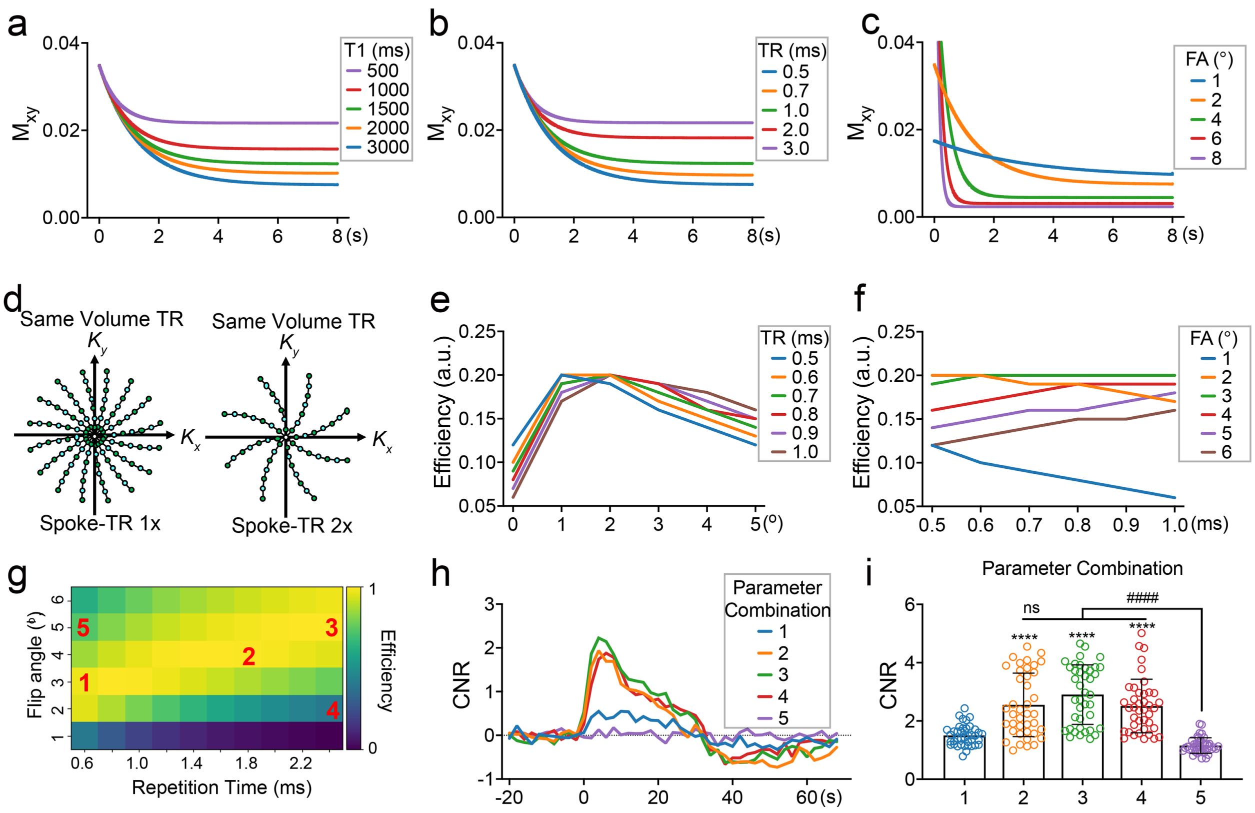


**Figure S4. TR and FA selections in SORDINO**. Bloch equation modeling is similar to that shown in Figure 1d but uses various T1 **(a)**, TR **(b)**, and FA **(c)**. The results indicate that a shorter T1, longer TR, and smaller FA result in a higher magnetization steady state, translating to a stronger SORDINO signal. **(d)** For fMRI applications, it is essential to maintain a short volume-TR. At the same volume-TR, doubling the spoke-TR reduces the number of spokes available to sample k-space by half. Therefore, a short spoke-TR strategy is critical for SORDINO fMRI. **(e-g)** We further model the efficiency of SORDINO to detect a 2% T1 change, which is empirically observed during activation due to tO2-induced T1 shortening. The efficiency calculation includes an additional square root of TR to account for the direct influence of TR on the number of sampled spokes. This modeling assumes a baseline tissue T1 of 1900 ms at 9.4T. The results are displayed for several TRs while varying FA in **(e)**, and for several FAs while varying TRs in **(f)**. **(g)** A heatmap illustrates the effects of TR and FA on SORDINO's efficiency in detecting a 2% T1 change. **(h-i)** Functional contrast was validated using task-based fMRI with a 30-second forepaw stimulation design using within-subject comparisons across five TR-FA combinations labeled in **(g)** (n = 8 subjects, 5 trials per parameter per subject; Linear Mixed Effects Model, $\beta = 1.15$, $P<0.0001$; *P*-values are from Tukey's HSD). While the empirical data generally follow the modeling trend, the discrepancy between the two may be attributed to $B_1$-inhomogeneity, which arises from higher acquisition bandwidths required for shorter TR sequences, as well as by discrepancies between the assumed and actual transit times in rats. Results are expressed as mean ± SEM. ****$p<0.0001$ compared with parameter combination 5; ####$p<0.0001$ compare with indicated groups. ns, not significant.

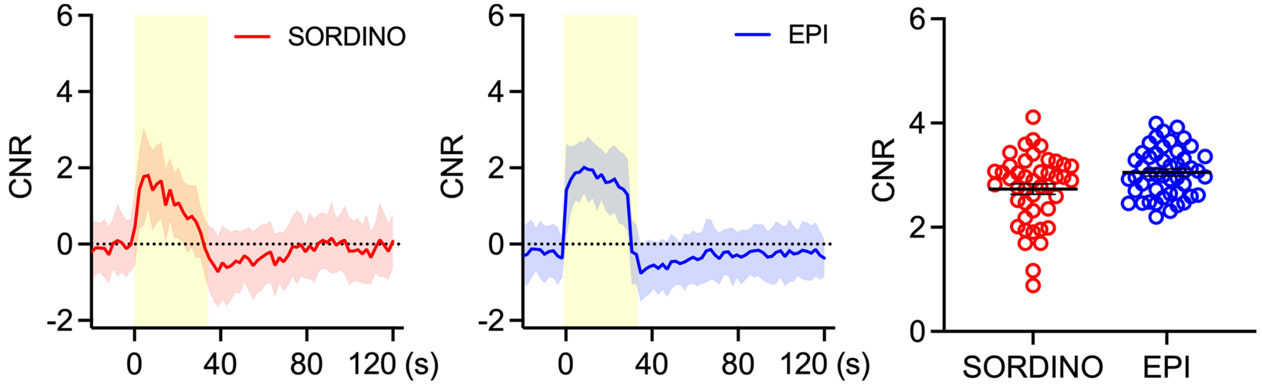


**Figure S5. SORDINO vs EPI Across Experimental Conditions.** SORDINO and EPI data were acquired with a temporal resolution of 2 seconds and a spatial resolution of 0.6 $mm^3$ (n = 4 subjects). The SORDINO dataset includes trials with free-breathing anesthetized rats, whereas all other data presented in the manuscript were in mechanically ventilated rats. SORDINO was acquired with: TR = 0.6006 – 1ms, flip angle = 3°, receiver bandwidth = 24 - 43 kHz, spokes= 1966 to 3330, matrix size = $44^3$, FOV = 26.4 $mm^3$. EPI parameters were: TR = 2s, TE = 14 ms, FA = 70°, receiver bandwidth = 250 kHz, matrix size = $44^2$, FOV = 26.4 $mm^2$, slices = 44 and slice thickness = 0.6 mm. SORDINO demonstrated comparable CNR to EPI, highlighting its robust performance across physiological states. Shaded regions are ± standard deviation; error bars are expressed as ± SEM.

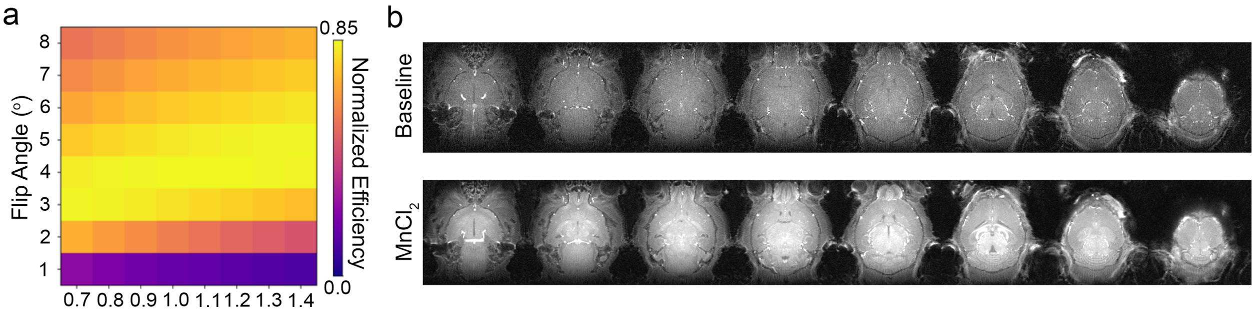


**Figure S6. SORDINO Efficiency with MEMRI at 9.4 T. (a)** To inform imaging parameters for MEMRI experiments with SORDINO, we modeled SORDINO efficiency using a 50% reduction in T1 from a baseline of 1900 ms. **(b)** After infusion of $MnCl_2$ with a minipump, SORDINO images of the mouse brain exhibited contrast in regions known to uptake manganese such as the pituitary gland, olfactory bulb, and cerebellum.

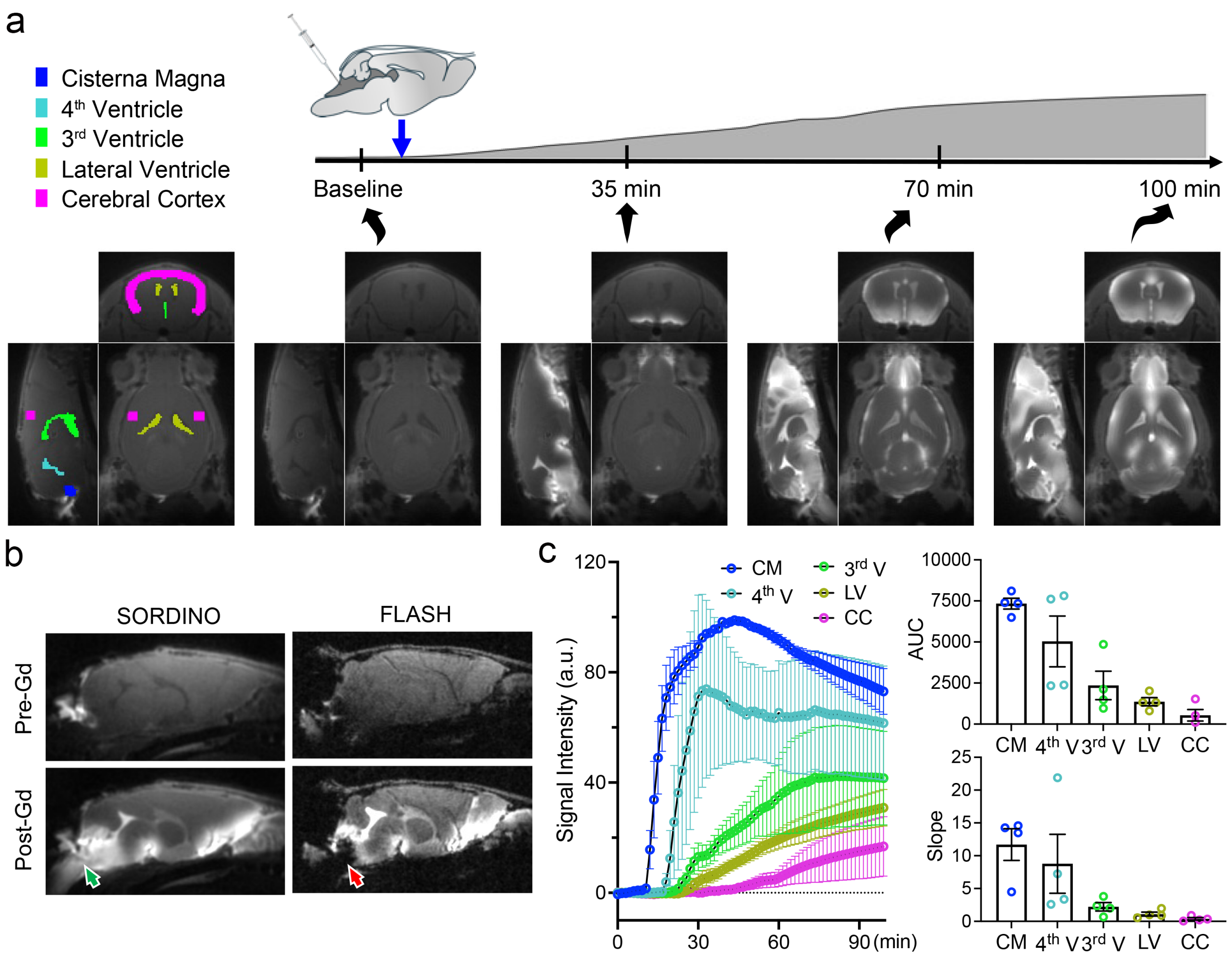


**Figure S7. Visualization of Glymphatic Dynamics using SORDINO. (a)** Representative regions of interest (ROIs) in key brain areas, including the cisterna magna (CM, blue), fourth ventricle (4th V, cyan), third ventricle (3rd V, green), lateral ventricle (LV, yellow), and cerebral cortex (CC, magenta), used to extract SORDINO signals over time. The Gd-enhanced signals are clearly visible in representative orthogonal brain views at baseline, 35 min, 70 min, and 100 min post-injection, illustrating the progressive distribution of Gd from CSF spaces to brain parenchyma. SORDINO data were acquired at 160 µm isotropic spatial resolution and 90 s temporal resolution. **(b)** Comparison of pre- and post-Gd signal intensity between SORDINO and FLASH sequence. SORDINO data show enhanced visualization of Gd contrast in the cisterna magna and adjacent regions with minimal susceptibility artifacts compared to FLASH. Green and red arrows highlight the region receiving direct Gd injection in SORDINO and FLASH, respectively. **(c)** Quantitative analysis of normalized signal intensity dynamics in the defined ROIs. Time-intensity curves demonstrate the temporal progression of Gd influx across ROIs, with the cisterna magna exhibiting the fastest and most pronounced signal changes. Bar plots summarize the area under the curve (AUC) and slope of Gd dynamics, with the injection site cisterna magna exhibiting the steepest uptake rate and greatest changes as expected. Data are presented as mean ± SEM, with individual data points shown (n = 5 mice). CM, cisterna magna; 4th V, fourth ventricle; 3rd V, third ventricle; LV, lateral ventricle; CC, cerebral cortex.

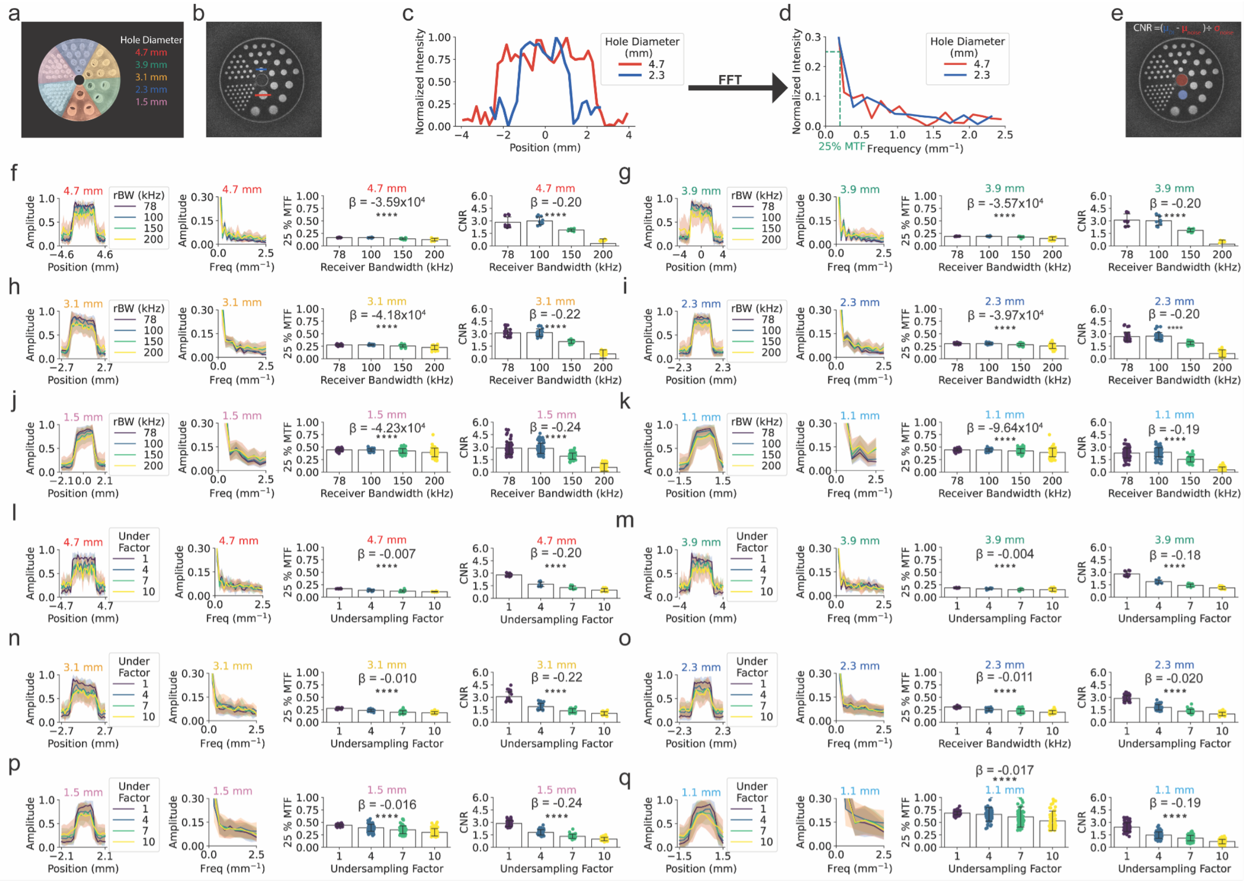


**Figure S8. Effect of Receiver Bandwidth and Undersampling Factor on SORDINO Spatial Resolution and Noise. (a)** A resolution phantom comprising holes of six different diameters (1.1 to 4.7 mm) filled with deionized water was used to estimate the effect of SORDINO imaging parameters on spatial resolution and noise. **(b,c)** As an illustrative example, line profiles were drawn through the center of the 4.7 (red) and 3.1 (blue) mm holes. To quantify the effect of imaging parameters on spatial resolution, we calculated the MTF which represents the imaging system's ability to preserve contrast across spatial frequencies, essentially serving as a frequency-domain characterization of imaging resolution. **(d)** The MTF is calculated by taking the FFT of line profiles drawn through the center of the phantom holes. To quantify spatial resolution, we chose an MTF threshold of 25%, which represents the spatial frequency at which 25% of contrast is preserved. **(e)** The effect of imaging parameters on noise was quantified by the CNR. As an illustrative example, the CNR was calculated by taking the difference between the phantom hole signal ROI (blue circle) and noise ROI (red circle) divided by the standard deviation of the noise ROI signal. In these experiments, we analyzed the number of line profiles or ROIs drawn over each phantom hole size as follows: 4.7 mm = 3, 3.9 mm = 3, 3.1 mm = 6, 2.3 mm = 10, 1.5 mm = 19, and 1.1 mm = 23, with each measurement repeated four times. Then, we assessed the effect of receiver bandwidth on SORDINO spatial resolution and noise. In these comparisons, we were particularly interested in whether the low receiver bandwidth strategy of SORDINO introduces spatial blurring via signal attenuation due to T2* decay during acquisition. **(a-k)** We found the 25 % MTF threshold decreases with increasing receiver bandwidth (linear mixed effects model, $\beta = -3.57$ to $-9.64 \times 10^{-4}$, $p<0.0001$). These results suggest that while receiver bandwidth has a small influence on SORDINO spatial resolution, any effect that does occur results in decreased spatial resolution at higher receiver bandwidths, likely due to increased artifacts and noise. **(a-k)** Similarly, we found that higher receiver bandwidths result in lower CNR (linear mixed effects model, $\beta = -0.017$ to $-0.024$, $p<0.0001$), likely due to increased noise variance at higher receiver bandwidths. Next, we assessed the impact of undersampling factor on SORDINO spatial resolution and noise. Like all MRI sequences, SORDINO exhibits a compromise between sufficient temporal (~ 2s) and spatial (~ 0.4 - 0.6 $mm^3$) resolution for rodent functional brain mapping. To balance this tradeoff, SORDINO data is undersampled in k-space which may introduce noise and artifacts into the reconstructed image due to violation of Nyquist's sampling criterion. **(l-q)** We found that higher undersampling factors reduced the 25% MTF threshold (linear mixed effects model, $\beta = -0.0040$ to $-0.017$, $p<0.0001$) with larger effects for smaller hole sizes. **(l-q)** Increasing undersampling factor reduces CNR (linear mixed effects model, $\beta$ = -0.18 to -0.23, $p<0.0001$). Together, these results suggest that careful consideration is needed when undersampling k-space significantly, especially when the goal is to resolve small ROIs. Results are expressed as mean ± SD. ****$p<0.0001$.

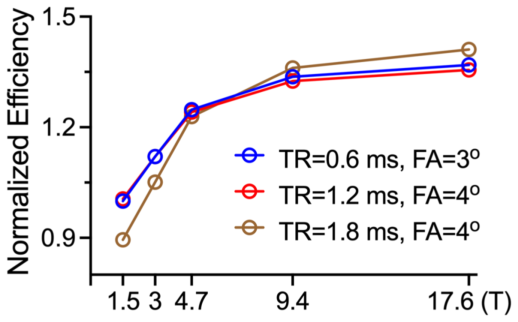


**Figure S9. SORDINO Efficiency to Changes in Tissue Oxygen as a Function of Field Strength.** Assuming baseline T1 values of 1035, 1272, 1600, 1900 and 2030 ms at field strengths of 1.5, 3, 4.7, 9.4, 17.6 T, respectively, and a 30 µM increase in $tO_2$ with a spin-lattice relaxivity (r1) of 0.3 $mM^{-1}s^{-1}$, SORDINO efficiency is expected to increase with field strength. Assuming baseline T1 values of 1035, 1272, 1600, 1900 and 2030 ms at field strengths of 1.5, 3, 4.7, 9.4, 17.6 T, respectively, and a30 µM increase in $tO_2$ with a spin-lattice relaxivity (r1) of 0.3 $mM^{-1}s^{-1}$, SORDINO efficiency is expected to increase with field strength. This is because the relationship between T1 and molecular oxygen is non-linear, as described by: $\frac{1}{T1_{new}} = \frac{1}{T1_{baseline}} = r_1.[\mathrm{tO2}\,]$. Since the additive increase in relaxation rate causes a proportionally larger drop in T1 when T1 is high, the efficiency of SORDINO improves at higher field strengths.
